# Proteomics Reveals Extracellular Matrix Injury in the Glomeruli and Tubulointerstitium of Kidney Allografts with Early Antibody-Mediated Rejection

**DOI:** 10.1101/2020.03.03.975672

**Authors:** Sergi Clotet-Freixas, Caitriona M. McEvoy, Ihor Batruch, Julie Van, Chiara Pastrello, Max Kotlyar, Madhurangi Arambewela, Alex Boshart, Sofia Farkona, Yun Niu, Yanhong Li, Olusegun Famure, Andrea Bozovic, Vathany Kulasingam, Peixuen Chen, Joseph Kim, Emilie Chan, Sajad Moshkelgosha, Tereza Martinu, Stephen Juvet, Igor Jurisica, Andrzej Chruscinski, Rohan John, Ana Konvalinka

## Abstract

Antibody-mediated rejection (AMR) accounts for >50% of kidney allograft losses. AMR is caused by donor-specific antibodies (DSA) against HLA and non-HLA antigens in the glomeruli and the tubulointerstitium, which together with inflammatory cytokines such as tumor necrosis factor alpha (TNFα) and interferon gamma (IFNɣ), trigger graft injury. Unfortunately, the mechanisms governing cell-specific injury in AMR remain unclear. We studied 30 for-cause kidney biopsies with early AMR, acute cellular rejection or acute tubular necrosis (‘non-AMR’). We laser-captured and microdissected glomeruli and tubulointerstitium, and subjected them to unbiased proteome analysis. 120/2026 glomerular and 180/2399 tubulointerstitial proteins were significantly differentially expressed in AMR vs. non-AMR biopsies (P<0.05). Basement membrane and extracellular matrix (ECM) proteins were significantly decreased in both AMR compartments. We verified decreased glomerular and tubulointerstitial LAMC1 expression, and decreased glomerular NPHS1 and PTPRO expression in AMR. Cathepsin-V (CTSV) was predicted to cleave ECM-proteins in the AMR glomeruli, and CTSL, CTSS and LGMN in the tubulointerstitium. We identified galectin-1, an immunomodulatory protein upregulated in the AMR glomeruli and linked to the ECM. Anti-HLA class-I antibodies significantly increased CTSV expression, and galectin-1 expression and secretion, in human glomerular endothelial cells. Glutathione S-transferase omega-1 (GSTO1), an ECM-modifying enzyme, was significantly increased in the AMR tubulointerstitium, and in TNFα-treated proximal tubular epithelial cells. IFNɣ and TNFα significantly increased CTSS and LGMN expression in these cells. Basement membranes are often remodeled in chronic AMR, and we demonstrated that this remodeling begins early in glomeruli and tubulointerstitium. Targeting ECM-remodeling in AMR may represent a new therapeutic opportunity.

**SIGNIFICANCE STATEMENT:** Antibody-mediated rejection (AMR) accounts for >50% of kidney allograft loss, and is caused by donor-specific antibodies against HLA antigens, which induce maladaptive responses in the kidney glomeruli and tubulointerstitium. This is the first unbiased proteomics analysis of laser-captured/microdissected glomeruli and tubulointerstitium from 30 indication kidney biopsies with early AMR, acute cellular rejection or acute tubular necrosis. >2,000 proteins were quantified in each compartment. We discovered that basement membrane and extracellular matrix (ECM) proteins were significantly decreased in both AMR compartments. Two ECM-modifying proteins, LGALS1 and GSTO1, were significantly increased in glomeruli and tubulointerstitium, respectively. LGALS1 and GSTO1 were upregulated by anti-HLA antibodies or AMR-related cytokines in primary kidney cells, and may represent therapeutic targets to ameliorate ECM-remodeling in AMR.

## INTRODUCTION

Kidney transplantation is the optimal treatment for end-stage kidney disease, but most grafts fail prematurely. Antibody-mediated rejection (AMR) is the leading cause of kidney graft failure, accounting for >50% of graft loss.^1^ AMR is characterized by graft injury (acute tubular necrosis (ATN) or thrombotic microangiopathy), donor-specific antibodies (DSA), and endothelial activation, as evidenced by C4d deposition in peritubular capillaries, microvascular inflammation, or increased endothelial injury markers. ATN indicates tubular cell injury and is a component of AMR.^2^ Chronic AMR is associated with an aggressive lesion, transplant glomerulopathy (TG), which portends poor prognosis.^3^ Multilayering of peritubular capillary basement membranes (BMs) and duplication of glomerular BM differentiate TG from other forms of AMR; these abnormalities likely arise from repeated cycles of endothelial cell injury. Caveolae formation in microvascular endothelial cells^4^ and development of anti-BM antibodies have been observed,^5–7^ but their significance is unclear. It is evident from the histopathological changes that injury to microvascular endothelial cells and BMs represent key sites of AMR-induced injury, initiated mainly by anti-HLA DSA.

Anti-HLA class-I (α-HLA-I) and class-II (α-HLA-II) antibodies exert pathogenic effects in human glomerular microvascular endothelial (HGMEC) and other cells.^8, 9^ Endothelial cells express HLA class-I constitutively, and HLA class-II in injury settings (e.g. exposure to interferon gamma (IFNɣ)).^10–12^ Ligation of HLA with α-HLA-I induces cell proliferation and activation of ERK and mTOR.^12–14^ These effects are enhanced by IFNɣ and tumor necrosis factor alpha (TNFα).^12^ Endothelial cells stimulated with α-HLA-I increased cytokine expression (e.g. IL-8 and CCL2).^15^ Importantly, α-HLA-I-activated endothelial cells display cytoskeletal alterations and interact with leukocytes.^16–19^ α-HLA-II induces necrosis in IFNɣ-treated endothelial cells,^10^ and exerts prothrombotic effects in HGMECs.^9^

The role of tubular cells in AMR is unclear. These cells express HLA and other immuno-modulatory proteins and interact with immune cells.^20, 21^ Proximal tubular epithelial cells (PTECs) increase inflammatory cytokine (IL-6, CCL2, and CXCL10) secretion in response to α-HLA-I, IFNɣ and TNFα.^22^ Both IFNɣ and TNFα accentuate antibody-mediated injury in AMR.^23–25^ Moreover, tubular proteins can be targeted by non-HLA antibodies,^22, 26^ which reinforces the rationale for studying AMR in the tubulointerstitium.

Identification of mechanisms underpinning cell-specific maladaptive responses in AMR may uncover new therapeutic targets. Transcriptome studies show that graft injury in AMR correlates with gene expression alterations.^27^ However, compartment-specific molecular alterations, or changes in proteome composition/expression are unknown. To address this gap, we conducted a discovery-based proteomics study in for-cause kidney biopsies. We selected 7 biopsies with pure, early AMR, and compared them to 23 graft age-matched ‘non-AMR’ biopsies with pure acute cellular rejection (ACR) or ATN. We isolated glomeruli and tubulointerstitium by laser-capture microdissection, and subjected them to mass-spectrometry (MS)-based proteome analysis. Working with ultra-small protein amounts, we identified >2000 proteins in each compartment. We demonstrated, for the first time, that BM and extracellular matrix (ECM) proteins were significantly decreased in AMR, in both compartments. We identified galectin-1 (LGALS1), an ECM-related immunomodulatory protein increased in AMR glomeruli and glutathione S-transferase omega-1 (GSTO1), an ECM-modifying enzyme increased in the AMR tubulointerstitium. These proteins represent potential therapeutic targets to prevent ECM-remodeling in AMR.

## METHODS

### Study design and patient population

We carefully selected early, for-cause kidney allograft biopsies with a diagnosis of acute AMR, and graft age-matched cases with ACR or ATN. Patients were identified by searching the CoReTRIS registry for the period of 2008-2015.^28^ Inclusion criteria encompassed a kidney transplant with an allograft biopsy within the first 3 months post-transplantation, with a diagnosis of pure AMR, ACR or ATN. Exclusion criteria were a diagnosis of mixed rejection or insufficient formalin-fixed paraffin-embedded biopsy material to perform laser-capture microdissection. For each patient, different sections from the same 18G-needle biopsy core were employed for histopathology evaluation, proteome analysis, and orthogonal verification studies. All biopsies were assessed by a renal pathologist (R.J.) and scored according to the Banff classification (2017).^29^ Anti-HLA antibodies were assessed using a Luminex single-antigen bead assay and adjudicated by the local HLA laboratory, as part of clinical standard of care.

## Histology

### Microscopic Pathology

From the formalin-fixed, paraffin-embedded biopsy material of our study cases, sections were cut at 3μm, subjected to PAS, PASM, Trichrome and H&E stains, and examined by light microscopy. C4d was detected on frozen sections by immunofluorescence. Morphologic features were diagnosed and scored according to the updated Banff classification^29^. For electron microscopy, tissue was examined following appropriate fixation using a JEOL-100S microscope. Glomerular basement membrane thickness was examined by standard methodology on 2 glomeruli per biopsy. Approximately 10-15 capillary loops per glomerulus and 10-15 peritubular capillaries per biopsy were studied at 8000x magnification. Subendothelial new basement membrane formation was reported as the proportion of loops with any degree of new basement membrane formation. Peritubular capillary basement membrane multilayering, as well as glomerular and peritubular capillary endothelial cell swelling were semiquantitatively assessed on a scale of 0 to 3. Scale legend: 0 = none; 1 = mild (<25%); 2 = moderate (25-50%); 3 = severe (>50%).

### Immunostaining

For LAMC1 immunohistochemistry, antigen retrieval was performed in 5μm sections by heating the samples with 0.01M Na-citrate (pH6) in a pressure cooker for 5 minutes. Sections were then incubated with rabbit anti-LAMC1 antibody (dilution 1:100, HPA001909, Atlas Antibodies). HRP-conjugated anti-rabbit (BA-1000, Vector Labs) was used as secondary antibody. Binding of antibodies was detected using the Liquid DAB + Substrate Chromogen System (Dako). Samples were counterstained with hematoxylin to visualize nuclei, and slides were digitally scanned in a ZEISS Axio Scan.Z1 system. To assess protein expression, glomerular and tubulointerstitial areas were differentially outlined, and the intensity of the positive pixels was quantified using the ImageScope software (version 12.3.2.8013).

We also assessed glomerular expression of NPHS1 and PTPRO by immunofluorescence. Deparaffinised sections were heated at pH9 to retrieve antigens, and incubated with rabbit anti-NPHS1 antibody (dilution 1:1,000, ab216341, Abcam), and with mouse anti-PTPRO (dilution 1:50, MABS1221 Millipore). Cy3-conjugated anti-rabbit IgG (A11035, Invitrogen) and FITC-conjugated anti-mouse IgG (A11029, Life Technologies) were used as secondary antibodies to detect NPHS1 and PTPRO, respectively. DAPI staining was employed to visualize the nuclei, and slides were digitally scanned in a ZEISS Axio Scan.Z1 system. To assess protein expression, glomerular areas were selected, and positive pixels were quantified using a ZEISS Imaging Software (ZEN 2 blue edition). For our immunostaining studies, data were expressed as mean intensity of positive pixels, and normalized to the μm^2^ of area analyzed.

### Proteomics

#### Laser-capture microdissection and sample preparation for proteomics analysis

For each study case, remaining tissue from formalin-fixed paraffin-embedded biopsy was sectioned at 8μm, and stained with hematoxylin. Using a laser-capture microdissection system (Leica), the glomeruli and tubulointerstitium were then isolated and collected in a tube with 35μL of protein extraction buffer containing 10mM Tris, 1mM EDTA, and 0.002% Zwittergent in MS-grade pure water (Sigma). To obtain comparable protein amounts across all samples, in each biopsy section, we standardized the amount of captured glomeruli and tubulointerstitium to 350,000μm^2^. In each biopsy section, this area included an average of 23 captured glomeruli (glomerular compartment) or 4-5 captured tubulointerstitial areas (tubulointerstitial compartment).The total protein amount per sample was below the limit of detection of standard protein quantification assays. Samples were vortexed for 2 minutes and centrifuged at 12,000G for 2min. To reverse formaldehyde crosslinking, samples were heated for 90min at 98°C, being vortexed every 15min. Samples were then centrifuged at 12,000G for 2min and subjected to sonication for 1h. Following another centrifugation at 12,000G for 2min, proteins were then digested into peptides with 0.5μg of MS grade trypsin (Promega) diluted in 50mM ammonium bicarbonate (NH₄HCO₃), overnight and at 37°C. After tryptic digestion, peptides were reduced by adding 2μL of 100mM dithiothreitol in NH₄HCO₃ (final concentration 5.13mM) and heating the samples at 95°C for 5min. Samples were then acidified by addition of 10μL of 0.5% v/v trifluoroacetic acid + 0.15% v/v formic acid in MS-grade pure water (Sigma). Peptides were extracted and desalted with 10μL OMIX C18 MB tips (Agilent), eluted in 3μL of 65% v/v acetonitrile, and diluted to 41μL with 0.1% v/v formic acid in mass spectrometry (MS)-grade pure water.

#### Tandem mass spectrometry (MS/MS)

For each study biopsy section, the glomerular and tubulointerstitial fractions were randomized and subjected to MS on a Thermo Scientific EASY-nLC1000 system, coupled to a Q-Exactive Plus hybrid quadrupole-orbitrap mass spectrometer using a nano-electrospray ionization source (Thermo Scientific). Samples were run on a 60-minute gradient of increasing concentrations of Buffer B (100% acetonitrile) in 0.1% formic acid/99.9% MS-grade water (Thermo Scientific). The method started at 1% Buffer B, and the concentration was increased to 5% at 2min, with subsequent increases to 35% (49min), 65% (52min) and 100% (53min). For each sample, 18μL of eluted peptides were injected onto a 3.3cm C18 pre-analytical column (IntegraFrit capillary, New Objective; inner diameter: 75μm; bead size: 5μm; Agilent Technologies) and then passed through a C18 resolving analytical column (PicoTip emitter, inner diameter: 15cm x 75μm; tip: 8μm tip; bead size: 3μm; Agilent Technologies). The spectra were obtained under data-dependent acquisition mode, consisting of full MS1 scans (m/z range: 400-1500; resolution: 70,000) followed by MS2 scans of the top 15 parent ions (resolution: 17,500).

#### Protein identification and quantification

For protein identification, the RAW files of each MS run were generated by XCalibur software v3.0.63 (Thermo Scientific). Raw data were analyzed by MaxQuant software (version 1.5.3.28) and searched in Andromeda against the human Uniprot FASTA database (HUMAN5640_sProt– 072016, update of July 20, 2016). Proteins and peptides were identified with a false discovery rate of 1%. For peptide identification, a minimum length of 6 amino acids was selected. The false positive rate was determined using reversed mode. Trypsin/P was selected as digestion enzyme, and a maximum of 2 missed cleavages was enabled. While cysteine carbamidomethylation was selected as a fixed modification, methionine oxidation and N-terminal acetylation were set as variable modifications. The initial peptide tolerance against a ‘human-first-search’ database was set to 20ppm. The main search peptide mass tolerance was 40ppm, and the fragment mass MS/MS tolerance was 0.5Da. Matching between runs was selected. Normalized label-free quantification (LFQ) of proteins was derived from extracted ion current information from razor and unique peptides with a minimum ratio count of 1. The mass spectrometry proteomics data have been deposited to the ProteomeXchange Consortium via the PRIDE partner repository^30^ with the dataset identifier PXD017580.

Proteomics data was analyzed using Perseus software (version_1.5.6.0). Reverse hits and contaminants were manually checked and removed. We examined the distribution of log2 transformed LFQ intensity values of all proteins quantified in each sample (**Fig.S1**). In the glomerular compartment, 28 of the 30 samples had protein intensity values that followed normal distribution. One ACR and one ATN case were excluded due to poor protein recovery, and non-normal distribution (**Fig.S1A**). In turn, the protein intensity values of all 30 tubulointerstitial fractions followed a normal distribution (**Fig.S1B**). We then subjected the zero-value intensities to imputation (assuming that low abundance values were missing), keeping a normal distribution, with a downshift of 1.8 standard deviations, and a width of 0.3. After imputation, we determined the differentially expressed proteins between the AMR and the non-AMR groups in glomeruli and tubulointerstitium, by comparing their mean log2 transformed LFQ intensities using the two-tailed independent t-test (P<0.05), followed by Benjamini-Hochberg adjustment (Q<0.2). All proteins selected for validation were identified by at least 5 unique peptides and present in all biopsy samples, except PTPRO, which was identified in 27/28 glomerular samples.

### Bioinformatics analyses

#### Principal component analysis

Principal component analysis (PCA) of the proteins differentially expressed in AMR in each compartment (120 in the glomeruli and 180 in the tubulointerstitium) was performed and plotted using ‘ggbiplot’ (Version 0.55) and RStudio (Version 1.1.463) in R (Version 3.6.1).^31^ Twenty-eight samples were analysed in the glomerular compartment (7 AMR, 10 ACR, and 11 ATN cases) and 30 in the tubulointerstitial compartment (7 AMR, 11 ACR, and 12 ATN cases). For each sample, the log2 transformed LFQ intensity value of each protein was used for the PCA.

#### Protein expression correlation analysis

For all the differentially expressed proteins in each compartment (120 in the glomeruli and 180 in the tubulointerstitium), the Pearson correlation between the expression levels of two proteins was calculated, considering all possible combinations of two proteins. The correlation coefficients (r) were then represented in a correlation heatmap. For the associations of interest, the significance of the correlation was calculated using Graphpad Prism 8.0. P<0.05 was considered statistically significant.

#### Gene ontology and pathway enrichment analysis

Gene ontology terms enriched among the 120 proteins differentially expressed in the AMR vs. non-AMR glomeruli, and among the 180 proteins differentially expressed in the AMR vs. non-AMR tubulointerstitium, were identified using the BiNGO plug-in in Cytoscape (version 3.4.0). Terms related to biological processes, molecular function, and cell compartment were considered in the analysis. Similarly, pathDIP^32^ database 3.0 (http://ophid.utoronto.ca/pathDIP) was used for pathway enrichment analysis. For the statistical analyses in BiNGO and pathDIP, hypergeometric test with Benjamini-Hochberg correction was used, and Q<0.05 was considered significant.

The significantly enriched pathways emerging from PathDIP analysis in each compartment were classified using a reduction algorithm. Through this algorithm, pathway categories were mapped to high-level pathway terms in two ontologies: PathwayOntology (https://bioportal.bioontology.org/ontologies/PW) and Reactome^33^ (https://reactome.org/). First, each pathway category was mapped to ontology terms based on the similarity of pathway names and ontology terms using Python (version 3.6.8). Next, each pathway category was mapped to an ancestor of its associated ontology term, at the first (Reactome) or second (PathwayOntology) level of the ontology. When there were multiple ancestors, the best one was manually selected. Ancestor terms not pertinent to this work were grouped in the category “Other”.

#### Protein-protein interaction and network analysis

Physical protein-protein interactions between the 300 proteins altered in AMR were collected using the Integrated Interactions Database^34^ (IID, version 2018-11). Interactions experimentally validated or predicted, and annotated as present in the human kidney were retained and visualized using NAViGaTOR 3.0.13^35^ (http://ophid.utoronto.ca/navigator). The color of the nodes reflects the category to which each protein was computationally assigned. If a protein was annotated with pathways from multiple categories, only the category with the highest number of pathways (“other” excluded) was used for annotation.

#### Analysis of external dataset

We examined a publicly-available transcriptome dataset (GSE36059).^27^ We identified all the GSM files corresponding to the microarray gene expression data from control (n=281) and AMR (n=65) kidney biopsies, and downloaded them from the Gene Expression Omnibus public functional genomics data repository (https://www.ncbi.nlm.nih.gov/geo/). We then conducted a differential gene expression analysis of AMR vs control in R software (version 3.5.2), using a limma moderated t-test followed by Benjamini-Hochberg adjustment (Q<0.05). We cross-referenced our list of proteins differentially expressed in the AMR vs non-AMR glomeruli and tubulointerstitium with the 1,275 genes significantly differentially expressed in AMR vs control biopsies, and studied the overlap between data sets.

#### Protease prediction analysis

MEROPS^36^ database (https://www.ebi.ac.uk/merops/) was used to identify proteases predicted to cleave the ECM-related proteins in each AMR compartment. Briefly, the UniProt accession numbers corresponding to the 62 ECM-related proteins differentially expressed in the AMR vs. non-AMR glomeruli, and the 66 ECM-related proteins downregulated in the AMR vs. non-AMR tubulointerstiitum, were evaluated in MEROPS to identify cleavage sites by known proteases. Proteases were ranked manually in descending order, according to the number of targets in our data they were predicted to cleave.

### Cell culture

#### Human glomerular microvascular endothelial cells

Primary human glomerular microvascular endothelial cells (HGMECs, Cell Systems) were cultured in Endothelial Cell Growth Media MV (Promocell), supplemented with a ready-to-use kit containing 5% v/v dialyzed fetal calf serum (FCS), 10ng/mL epidermal growth factor (EGF), 1μg/mL hydrocortisone, and 90μg/mL heparin. Fifty units/mL penicillin and 50g/mL streptomycin were added. All the experiments were performed at passage 5. After reaching ∼70% confluence, cells were serum-starved for 16h in media containing 0.5% FCS. Cells were then treated with 1, 5, or 10μg/mL of mouse anti-human HLA class-I antibody (clone W6/32, ab23755, Abcam) for 7.5, 15, 30, 60, or 120min (signaling experiments), or for 18h or 24h (gene/protein expression studies). The same concentration of mouse IgG2α (ab18413, Abcam) was administered as isotype control, and PBS was used as vehicle. To study response to cytokines, HGMECs were also exposed to 10ng/µL of TNFα (Sigma) or 1,000U/mL of IFNγ (Sigma) for 24h. For these experiments, 0.1% bovine serum albumin (BSA) in PBS was used as vehicle. All treatments were prepared in FCS-free media. After stimulation, cells were washed with PBS, and harvested with 0.25% trypsin-EDTA (Life Technologies) for 5min at 37°C. Cell pellets were then snap-frozen and stored at −80°C until further analysis. Cell supernatants were also collected, cleared at 1,000G and 4°C for 10min to remove cell debris, and stored at −80°C until further analysis.

#### Human proximal tubular epithelial cells

Primary human proximal tubular epithelial cells (PTECs, Lonza) were cultured in Dulbecco’s modified Eagle’s medium (DMEM) containing 5.55mM D-glucose, 4mM L-glutamine, and 1mM sodium pyruvate, and supplemented with 10% v/v dialyzed fetal bovine serum (FBS), 10 ng/ml EGF, 1x of Transferrin/Insulin/Selenium (Invitrogen), 0.05M hydrocortisone, 50 units/ml penicillin, and 50g/ml streptomycin, as previously described.^37^ All the experiments were performed at passage 5. After reaching ∼70% confluence, cells were serum-starved for 16h and treated with 20ng/µL of TNFα, 1,000U/mL of IFNγ, or 1ng/µL of lipopolysaccharide (LPS) (all from Sigma) for 24h. 0.1% BSA in PBS was used as vehicle. At the end of the treatment, cells were washed with PBS and harvested with 0.25% trypsin-EDTA (Life Technologies) for 5min at 37°C. Cell pellets were then snap-frozen and stored at −80°C until further analysis.

### Gene expression

Total RNA was extracted from cell pellets using the RNAeasy Mini Kit (Qiagen). After quantifying the RNA concentration using Nanodrop (Thermo Scientific), 300ng of RNA were retrotranscribed to cDNA using the High Capacity cDNA Reverse Transcription Kit (Applied Biosystems). Gene levels of CHD5, PECAM1, VWF, ACTA2, CCL2, IL6, CXCL8, CXCL10, VCAM1, LGALS1, TAP1, HLA-C, CTSV, CTSL, CTSS, CASP1, LGMN, MMP2, MMP9, and GSTO1 were measured by real-time quantitative PCR using Power SYBR® Green PCR Master Mix (Applied Biosystems) in a StepOnePlus System (Applied Biosystems). Human lung fibroblasts (Hel299 cell line), used as positive control for ACTA2 gene expression, were kindly supplied by Terrance Ku (Humar laboratory, University Health Network). In each experiment, gene expression data were normalized to the most stable of four housekeeping genes, across conditions: ACTB, GAPDH, VCL or RPL31. All primer sequences are summarized in **Table S1**.

**Table 1.**
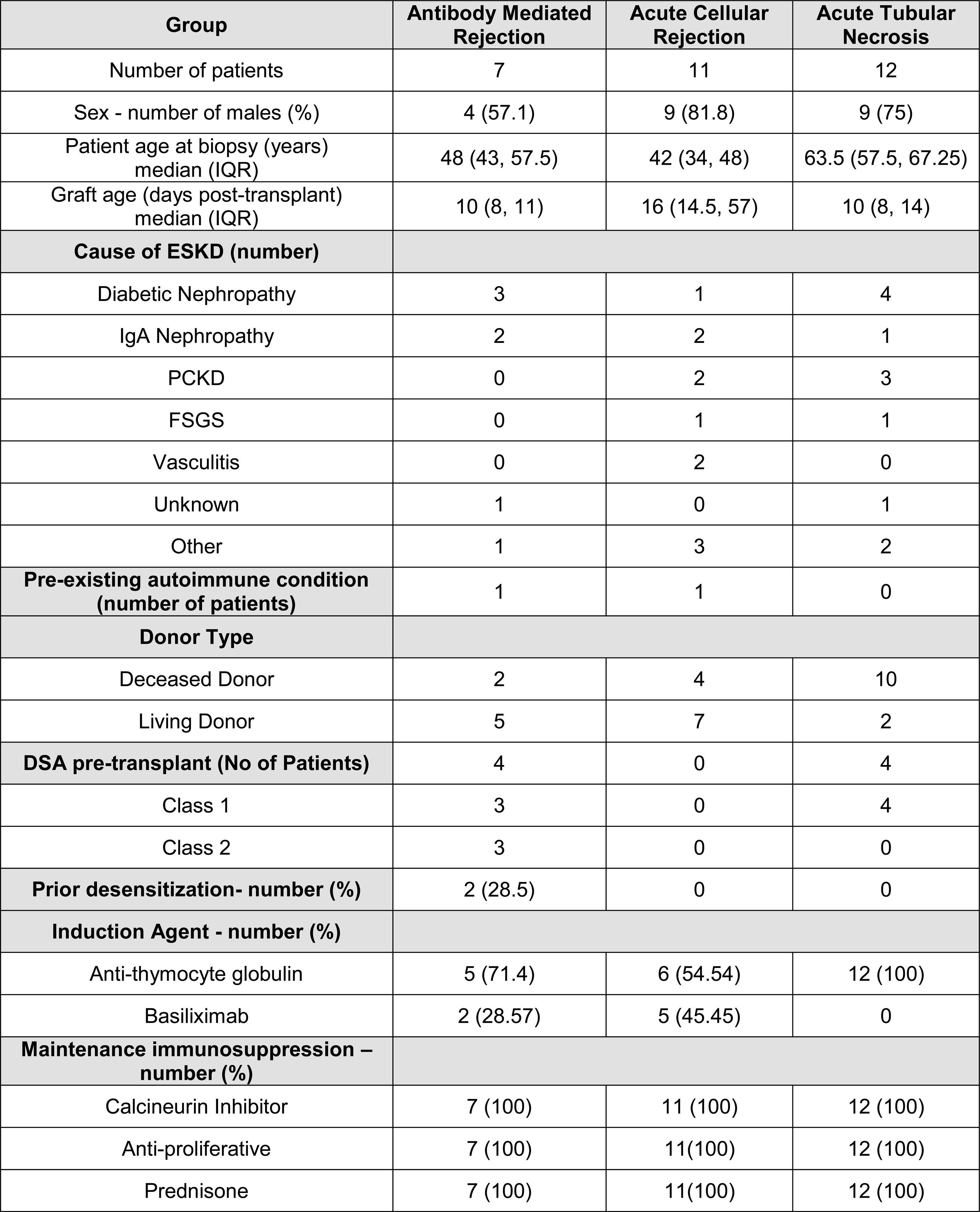
Clinical parameters of the patient cohort. IQR, interquartile range; ESKD, end-stage kidney disease; PCKD, polycystic kidney disease; FSGS focal segmental glomerulosclerosis; DSA, donor-specific antibodies.

| Group | Antibody Mediated Rejection | Acute Cellular Rejection | Acute Tubular Necrosis |
| --- | --- | --- | --- |
| Number of patients | 7 | 11 | 12 |
| Sex - number of males (%) | 4 (57.1) | 9 (81.8) | 9 (75) |
| Patient age at biopsy (years) median (IQR) | 48 (43, 57.5) | 42 (34, 48) | 63.5 (57.5, 67.25) |
| Graft age (days post-transplant) median (IQR) | 10 (8, 11) | 16 (14.5, 57) | 10 (8, 14) |
| **Cause of ESKD (number)** |  |  |  |
| Diabetic Nephropathy | 3 | 1 | 4 |
| IgA Nephropathy | 2 | 2 | 1 |
| PCKD | 0 | 2 | 3 |
| FSGS | 0 | 1 | 1 |
| Vasculitis | 0 | 2 | 0 |
| Unknown | 1 | 0 | 1 |
| Other | 1 | 3 | 2 |
| **Pre-existing autoimmune condition (number of patients)** | 1 | 1 | 0 |
| **Donor Type** |  |  |  |
| Deceased Donor | 2 | 4 | 10 |
| Living Donor | 5 | 7 | 2 |
| **DSA pre-transplant (No of Patients)** | 4 | 0 | 4 |
| Class 1 | 3 | 0 | 4 |
| Class 2 | 3 | 0 | 0 |
| **Prior desensitization- number (%)** | 2 (28.5) | 0 | 0 |
| **Induction Agent - number (%)** |  |  |  |
| Anti-thymocyte globulin | 5 (71.4) | 6 (54.54) | 12 (100) |
| Basiliximab | 2 (28.57) | 5 (45.45) | 0 |
| **Maintenance immunosuppression – number (%)** |  |  |  |
| Calcineurin Inhibitor | 7 (100) | 11 (100) | 12 (100) |
| Anti-proliferative | 7 (100) | 11(100) | 12 (100) |
| Prednisone | 7 (100) | 11(100) | 12 (100) |

### Protein expression

Cell pellets from HGMECs and PTECs were solubilized in lysis buffer (#9803, Cell Signaling), and total protein was extracted by mechanical homogenization. Protein concentration was determined using a Micro BCA protein assay kit (Thermo). To study the protein expression of HLA molecules in HGMECs by Western Blot, 10µg of protein were loaded onto 10% acrylamide gels, separated by SDS-PAGE, and transferred to a PVDF membrane (Millipore). Membranes were then blocked with 5% milk and incubated with mouse monoclonal anti-HLA class-I (1:4,000; ab23755, Abcam) or rabbit monoclonal anti-HLA class-II (1:3,000; ab157210, Abcam). For signaling experiments in HGMECs, blots were performed on 20µg of protein, and membranes were incubated with primary antibodies against phospho-p44/42 ERK (1:1,000; #9102, Cell Signaling) and total p44/42 ERK (1:2,000; #9102 Cell Signaling). To measure GSTO1 protein expression in PTECs, blots were performed on 40µg of protein, and membranes were incubated with 1/500 of anti-GSTO1 rabbit polyclonal antibody (HPA037603, Atlas Antibodies). Control for protein loading in HGMECs was performed by reblotting membranes using a rabbit polyclonal antibody for alpha-tubulin (ab176560, Abcam) or a mouse monoclonal antibody for GAPDH (CB1001, Sigma). The secondary antibodies were anti-rabbit (A0545, Sigma) and anti-mouse (P0447, Dako). Following detection in a Gel-Imaging System (Bio-rad), bands were quantified by densitometry using Image J software.

### Cytokine and galectin-1 secretion

Frozen aliquots of supernatants from HGMECs were thawed and subjected to a custom multiplex bead kit (R&D Systems) to assess the levels of IL-4, IL-8, and CCL2. Samples were prepared at a 1:1 dilution with assay buffer, as suggested by the manufacturer. Diluted samples and standards were run in duplicate. Biomarker concentrations were obtained using a Bio-Plex® MAGPIXTM Multiplex reader (Bio-Rad Laboratories, Hercules, CA). For all the analysed cytokines, any value falling below the lower limit of detection was assigned a concentration of 0ng/ml.

To determine the secreted levels of LGALS1, cell supernatants from HGMECs were analysed in duplicate using the colorimetric LGALS1 Human ELISA Kit (KA5065, Abnova). Supernatants were diluted 1:15 in sample diluent and added to the plate wells, which were pre-coated with an anti-LGALS1 capture antibody. After an incubation of 90min at 37°C, plates were washed with PBS and a biotinylated anti-LGALS1 detection antibody was added. Samples were then incubated for 60min at 37°C, and washed again with PBS. Detection was performed by adding an Avidin-Biotin-Peroxidase Complex (ABC-HRP) and incubating the plates with color developing reagent for 15min at 37°C. The emitted colorimetric signal was recorded at 450nm in a Cytation 5 plate reader (BioTek).

### Cellular metabolic function

Metabolic function was assessed in HGMECs and PTECs by measuring the extracellular acidification rate (ECAR, indicator of glycolysis) and oxygen consumption rate (OCR) in a Seahorse XFe96 analyzer (Agilent). At confluence, cells were detached with 0.25% trypsin-EDTA for 5min at 37°C, and subsequently seeded in a Seahorse XF96 Cell Culture Microplate at a density of 15,000 cells/well in 100μL of complete media. After letting them adhere for 4-6h, cells were serum-starved and exposed to the treatment of interest. One hour prior to the assay, starvation media was removed, and cells were washed with phenol-free basal media (Agilent) and exposed to 150μL of minimal substrate assay media (made by adding 2mM glutamine, 1mM pyruvate, and 5.55mM glucose to the basal media). The same treatment concentrations were maintained during this acclimatization step. During the Seahorse assay, ECAR and OCR were recorded at baseline and after metabolic stress. To induce metabolic stress, 25µL of oligomycin, p-trifluoromethoxy carbonyl cyanide phenyl hydrazone (FCCP), 2-deoxyglucose (2-DG), and Rotenone + Antimycin A (Rot+AA) were sequentially injected into the microplate wells. After optimization, the following working concentrations for each metabolic drug were stablished: oligomycin 1µM; FCCP 0.6µM for HGMECs and 0.3µM for PTECs, 2-DG 100mM; Rot 1µM; AA 1µM.

### Oxidative stress

Oxidative stress was assessed in HGMECs and PTECs by measuring the intracellular levels of superoxide ion with the Cellular ROS Assay Kit (Red) (Abcam) following the manufacturer instructions. At confluence, cells were detached with 0.25% trypsin-EDTA for 5min at 37°C, and subsequently seeded in black 96-well microplates at a density of 15,000 cells/well in 100μL of complete media. After letting the cells adhere for 4-6h, they were starved and exposed to the treatment of interest. After each treatment, starvation media was removed and cells were washed with PBS. Cells were stained with 100μL of ROS Red Working Solution for 45min at 37°C. Changes in fluorescence intensity were recorded at an Ex/Em = 520/605nm in a Cytation 5 plate reader (BioTek).

### Intracellular levels of DNA and ATP

Intracellular levels of DNA and ATP in HGMECs and PTECs were measured with a Cyquant Assay Kit (Thermo Scientific) and a CellTiter-Glo 2.0 Assay Kit (Promega), respectively, following manufacturer instructions. At confluence, cells were detached with 0.25% trypsin-EDTA for 5min at 37°C, and subsequently seeded in black (for DNA) or white (for ATP) 96-well microplates at a density of 15,000 cells/well in 100μL of complete media. After letting the cells adhere for 4-6h, they were starved and exposed to the treatment of interest. After each treatment, media was removed and cells were washed with PBS. To measure DNA levels, cells were exposed to 200µL of CyQUANT dye in cell-lysis buffer for 5min at RT, and fluorescence was recorded at an Ex/Em = 480/520nm in a Cytation 5 plate reader (BioTek). To measure ATP levels, cells were exposed to 100µL of CellTiter-Glo 2.0 Reagent and contents were mixed for 2min on an orbital shaker to induce cell lysis. Plates were then incubated at room temperature for 10min to stabilize the luminescent signal, which was recorded in a Cytation 5 plate reader (BioTek).

## Statistical Analyses

Normal distribution of each study variable was examined using the Shapiro-Wilks normality test. We assessed differences between groups using the independent t-test for variables following a normal distribution, and the Wilcoxon-Mann-Whitney non-parametric test for variables not following a normal distribution. Pearson correlation coefficient was calculated on log2-transformed LFQ protein intensity values. The Benjamini-Hochberg correction was used for multiple-hypothesis testing. P<0.05 was considered significant. Data are reported as mean ± standard error.

## RESULTS

### Study population

Seven patients with pure AMR were compared to 23 patients with other forms of non-AMR allograft injury, namely ACR and ATN (**Table 1**). Median graft age for all groups was ∼10 days, indicating that we were studying early cases. Four AMR patients had DSA prior to transplant. None of the ATN or ACR cases showed histopathologic signs of AMR (**Table 2**). ACR cases showed the highest scores for interstitial inflammation, tubulitis and total inflammation. The AMR cases showed microvascular inflammation, which was mostly absent in the other groups. There was no evidence of chronic injury in any group, including no evidence of TG (**Fig.S2A**). The lack of evident chronic lesions was confirmed by electron microscopy (**Fig.S2B**). There were no significant differences between groups in the glomerular BM thickness, (**Fig.S2B,C**), new BM formation (**Fig.S2D**), or endothelial cell swelling in the glomeruli (**Fig.S2E**) and peritubular capillaries (**Fig.S2F**).

**Table 2.**
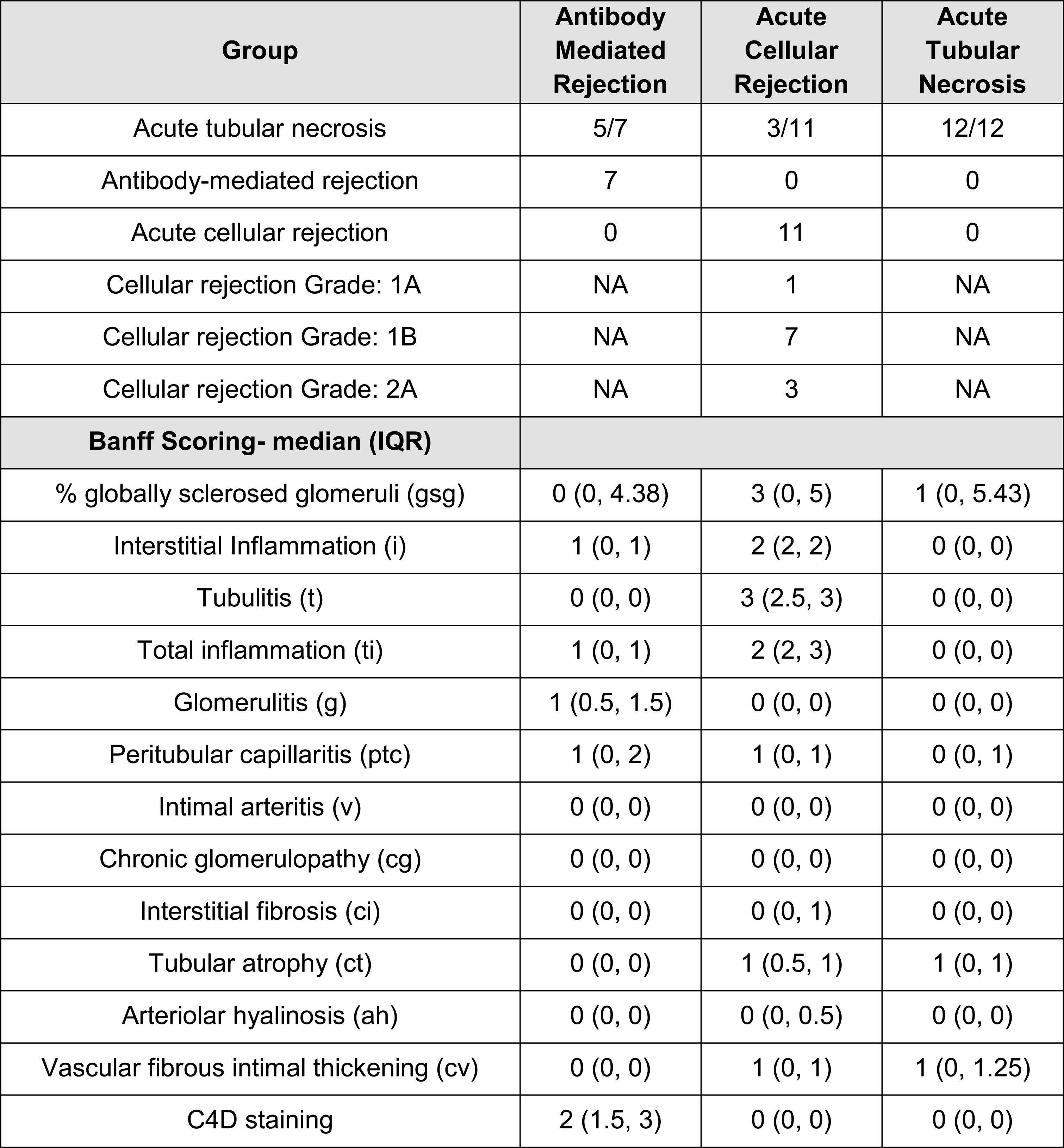
Kidney biopsy findings of the patient cohort. IQR, interquartile range; NA, not applicable.

| Group | Antibody Mediated Rejection | Acute Cellular Rejection | Acute Tubular Necrosis |
| --- | --- | --- | --- |
| Acute tubular necrosis | 5/7 | 3/11 | 12/12 |
| Antibody-mediated rejection | 7 | 0 | 0 |
| Acute cellular rejection | 0 | 11 | 0 |
| Cellular rejection Grade: 1A | NA | 1 | NA |
| Cellular rejection Grade: 1B | NA | 7 | NA |
| Cellular rejection Grade: 2A | NA | 3 | NA |
| **Banff Scoring- median (IQR)** |  |  |  |
| % globally sclerosed glomeruli (gsg) | 0 (0, 4.38) | 3 (0, 5) | 1 (0, 5.43) |
| Interstitial Inflammation (i) | 1 (0, 1) | 2 (2, 2) | 0 (0, 0) |
| Tubulitis (t) | 0 (0, 0) | 3 (2.5, 3) | 0 (0, 0) |
| Total inflammation (ti) | 1 (0, 1) | 2 (2, 3) | 0 (0, 0) |
| Glomerulitis (g) | 1 (0.5, 1.5) | 0 (0, 0) | 0 (0, 0) |
| Peritubular capillaritis (ptc) | 1 (0, 2) | 1 (0, 1) | 0 (0, 1) |
| Intimal arteritis (v) | 0 (0, 0) | 0 (0, 0) | 0 (0, 0) |
| Chronic glomerulopathy (cg) | 0 (0, 0) | 0 (0, 0) | 0 (0, 0) |
| Interstitial fibrosis (ci) | 0 (0, 0) | 0 (0, 1) | 0 (0, 0) |
| Tubular atrophy (ct) | 0 (0, 0) | 1 (0.5, 1) | 1 (0, 1) |
| Arteriolar hyalinosis (ah) | 0 (0, 0) | 0 (0, 0.5) | 0 (0, 0) |
| Vascular fibrous intimal thickening (cv) | 0 (0, 0) | 1 (0, 1) | 1 (0, 1.25) |
| C4D staining | 2 (1.5, 3) | 0 (0, 0) | 0 (0, 0) |

### Identification of compartment-specific proteome changes in kidney AMR

The proteomics workflow is shown in **Fig.1A**. We identified 2,026 proteins in the glomeruli, and 2,399 proteins in the tubulointerstitium (**Fig.1B**). Among the proteins quantified in >50% samples/group (1,299 in the glomeruli and 1,842 in the tubulointerstitium), we confirmed compartment-specific enrichment of expected proteins. Podocyte markers (e.g. NPHS1 and PODXL) and endothelial markers (e.g. PECAM1 and PDGFRB) were exclusively found in the glomeruli, whereas tubule-specific markers (e.g. LRP2, CUBN and UMOD) were found only in the tubulointerstitium (**Fig.1C**).

**Figure 1.**
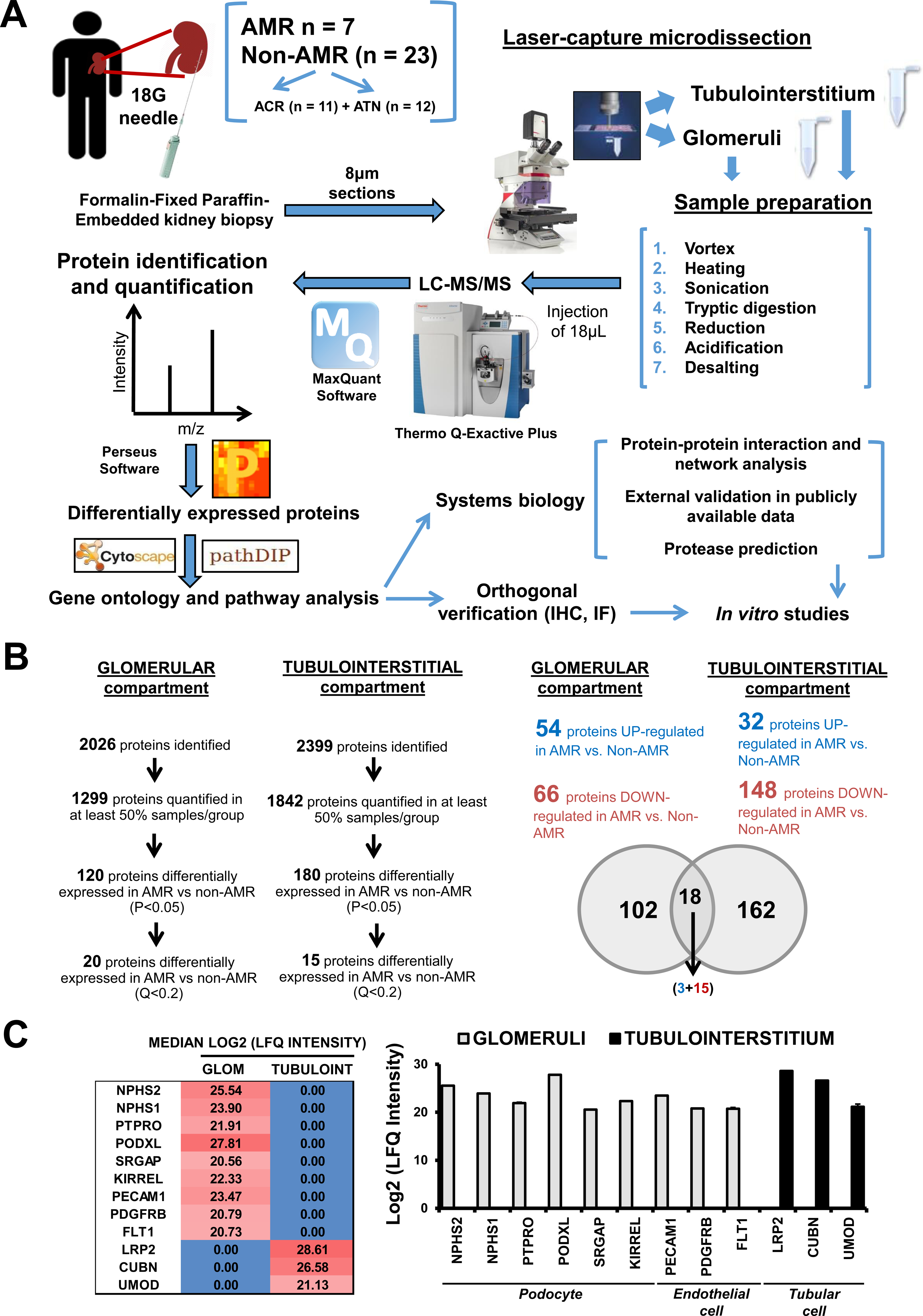
Experimental scheme. Panel A shows a simplified workflow, including laser-capture microdissection of glomerular and tubulointerstitial compartments, sample preparation followed by LC-MS/MS, data analysis by MaxQuant, statistical analysis to identify differentially expressed proteins (Perseus software), gene ontology term (Cytoscape software) and pathway enrichment analysis (pathDip portal and database), and downstream *in silico* analyses and verification/validation studies. Panel B illustrates the number of proteins identified and quantified, and the number of differentially expressed proteins between AMR and non-AMR biopsies, in each renal compartment. The Venn diagram illustrates the number of compartment-specific differentially expressed proteins between our AMR and non-AMR cases, as well as the overlap of significantly differentially expressed proteins between the glomerular and the tubulointerstitial compartments. Panel C shows compartment-specific enrichment of expected proteins. Whereas podocyte and endothelial cell markers were exclusively detected in glomerular fractions, tubule cell markers were only found in the tubulointerstitium. For each of the proteins of interest, the heatmap represents the median log2-transformed LFQ intensity across all samples, and in each compartment. AMR, antibody-mediated rejection; ACR, acute cellular rejection; ATN, acute tubular necrosis; LC-MS/MS, liquid chromatography followed by tandem mass spectrometry; IHC, immunohistochemistry; LFQ, label-free quantification; NPHS2, podocin; NPHS1, nephrin; PTPRO, receptor-type tyrosine-protein phosphatase O; PODXL, podocalyxin; SRGAP, SLIT-ROBO Rho GTPase-activating protein 2; KIRREL, Kin of IRRE-like protein; PECAM1, platelet endothelial cell adhesion molecule-1; PDGFRB, platelet-derived growth factor receptor beta; FLT1, vascular endothelial growth factor receptor-1; LRP2, megalin; CUBN, cubilin; UMOD, uromodulin.

We determined that 120 proteins were differentially expressed in the AMR compared to non-AMR glomeruli, and 180 proteins were differentially expressed in the tubulointerstitium. Eighteen of these proteins were differentially expressed in both compartments (**Fig.1C**). PCA of the intensity values of differentially expressed proteins confirmed a complete separation between AMR and non-AMR biopsies, in both compartments (**Fig.S3**).

Twenty proteins in the glomeruli and 15 in the tubulointerstitium were significantly altered between AMR and non-AMR after Benjamini-Hochberg adjustment (Q<0.2) (**Table S2**).

### AMR is associated with changes in ECM and BM proteins

We determined the dominant biological processes and pathways governing compartment-specific differences in AMR. In the glomeruli, gene ontology (GO) and pathway analysis demonstrated a significant enrichment of GO terms and signaling pathways that pinpointed 3 main protein groups. Forty-four proteins were related to immune system (**Fig.2A**, Table S3). The majority (e.g. TAP1, TAP2, and HLA-C) were increased in AMR compared to non-AMR glomeruli and participated in HLA class I-mediated antigen presentation. We also observed 62 proteins associated with cytoskeleton, cell adhesion, ECM and glomerular BM (**Fig.2B**, Table S3). Glomerular BM proteins, including nidogen (NID1), collagens (COL4A1, COL4A4), and laminin subunits (LAMC1, LAMA5, LAMB2), and podocyte-specific proteins NPHS1 and receptor-type tyrosine-protein phosphatase (PTPRO), were significantly decreased in AMR (**Fig.2B**). A third group involved 51 proteins related to metabolism and apoptosis (**Fig.2C**, Table S3). Proteins involved in metabolism were upregulated in AMR, whereas proteins cleaved by caspases during apoptosis were downregulated. Reassuringly, ‘unfolded protein response’ and ‘endoplasmic reticulum stress’ processes, which are intrinsically linked to HLA-mediated antigen presentation, were enriched among the proteins differentially expressed in the AMR glomeruli with Q<0.2 (**Table S3**).

**Figure 2.**
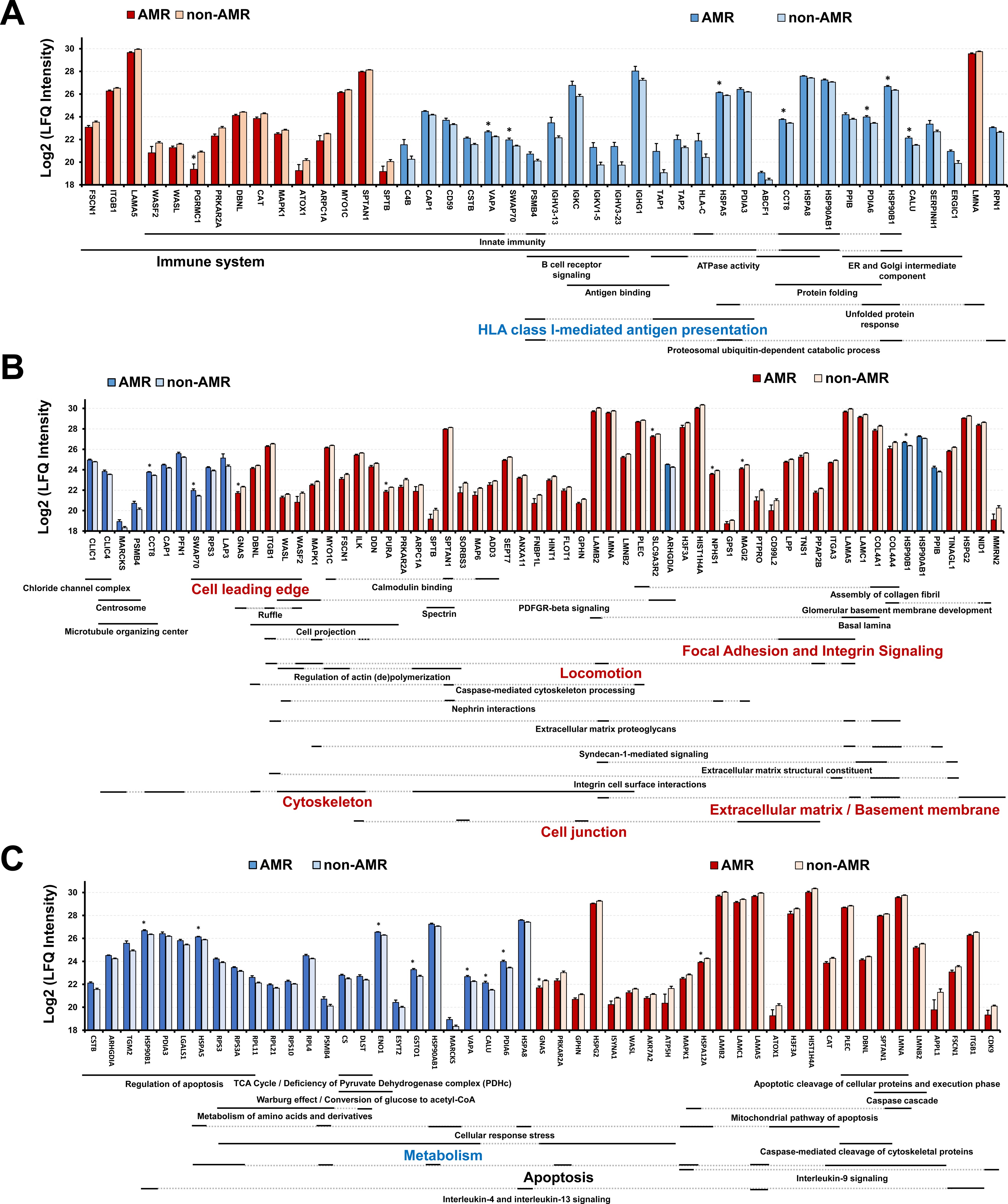
Significantly enriched gene ontology (GO) terms and molecular pathways among the 120 proteins differentially expressed in AMR compared to non-AMR glomeruli. The bar graphs illustrate the mean Log2-transformed ratios of the proteins belonging to each of the three main groups of proteins differentially expressed in the AMR vs non-AMR glomeruli (P<0.05): Immune system/ HLA class I-mediated antigen presentation (A), Extracellular matrix/ basement membrane/ cytoskeleton (B), and Metabolism/ apoptosis/ inflammation (C). The most representative significantly enriched GO terms and pathways are annotated below the names of the corresponding proteins. Blue colour indicates increase or enrichment in AMR, and red colour indicates decrease in AMR. Relative enrichment of GO terms was assessed using BinGO software in Cytoscape. Pathway enrichment was analysed using pathDIP. Data are expressed as mean ± SEM. *Q<0.2.

We also identified enriched GO terms and pathways in the AMR tubulointerstitium, where two prominent protein groups emerged. Sixty-six proteins decreased in the AMR tubulointerstitium were associated with cytoskeleton, ECM and BM (**Fig.3A**, Table S4). In contrast to glomeruli, we observed a higher representation of collagens and proteoglycans. Tubular BM components such as NID1, LAMA5, and LAMC1, downregulated in the AMR glomeruli, were also significantly decreased in the AMR tubulointerstitium. This enrichment for ECM-related processes and pathways in the AMR glomeruli remained evident among the proteins differentially expressed (Q<0.2) in the tubulointerstitium (**Table S4**). We identified a significant enrichment of metabolic processes and pathways in the AMR tubulointerstitium (**Fig.3B**, Table S4). Several proteins significantly increased in the AMR tubulointerstitium (e.g. CYP4A11, ETFA, and GSTO1) are enzymes with key roles in maintaining cellular redox homeostasis^38^. Unsurprisingly, ‘TNFα’ and ‘Interferon type-I signaling’ pathways were enriched among the 180 proteins differentially expressed in the AMR tubulointerstitium and their interactors, supporting the importance of TNFα and IFNɣ signaling in the AMR tubulointerstitium (**Table S5**).

**Figure 3.**
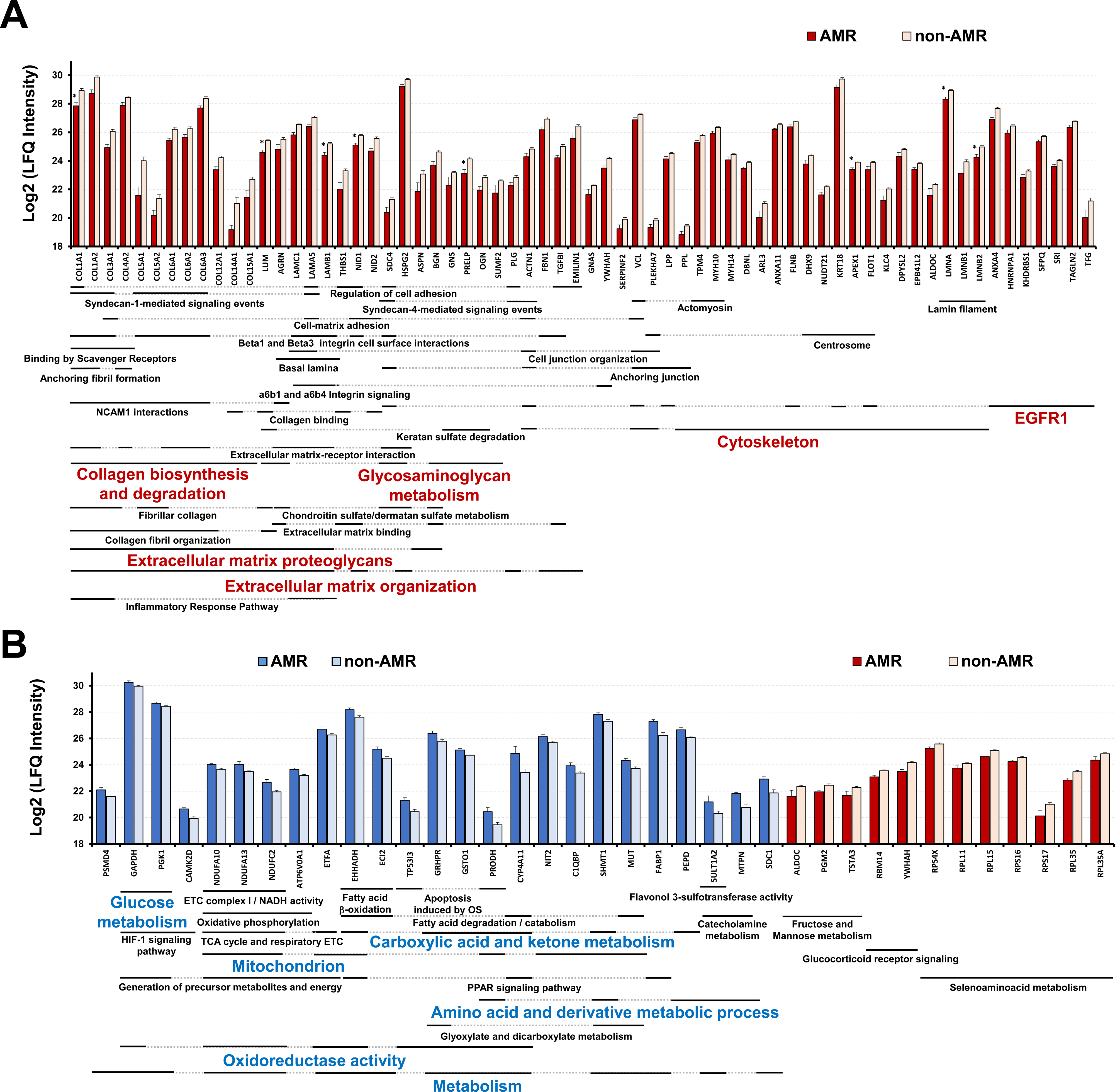
Significantly enriched gene ontology (GO) terms and molecular pathways among the 180 proteins differentially expressed in AMR compared to non-AMR tubulointerstitium. The bar graphs illustrate the mean Log2-transformed ratios of the proteins belonging to each of the two main groups of proteins significantly altered between AMR and non-AMR tubulointerstitium (P<0.05): Extracellular matrix / Glycosaminoglycans / Cytoskeleton (A) and Metabolism (B). The most representative significantly enriched terms and pathways are annotated below the names of the corresponding proteins. Blue colour indicates increase or enrichment in AMR, and red colour indicates decrease in AMR. Relative enrichment of GO terms was assessed using BinGO software in Cytoscape. Pathway enrichment was analysed using pathDIP. Data are expressed as mean ± SEM. *Q<0.2.

Using an unbiased approach, we confirmed the enrichment in ECM proteins in both AMR compartments. ‘ECM and cell communication’ was the main category among the pathways significantly enriched in AMR (**Fig.4A**, Table S6), and pathways falling into this category showed the highest significance (**Fig.4A**). Indeed, 45/300 proteins altered in AMR were assigned to ‘ECM and cell communication’; all 45 interacted with one another, or with other proteins altered in AMR (**Fig.4B**). Two proteins, LGALS1 and GSTO1, were significantly increased in AMR glomeruli and tubulointerstitium, respectively, and were highly connected to ECM and metabolic proteins (**Fig.4B**). Furthermore, we demonstrated significant co-expression of ECM proteins. In the glomeruli, LAMC1 correlated directly and strongly with NID1, LAMA5, and LAMB2. In the tubulointerstitium, LAMC1 expression correlated with NID1, LAMA5, LMNA, LMNB2, PRELP, and COL3A1 expression (**Fig.4C**, **Table S7**). Altogether, computational analyses pointed to ECM and BM as the key alterations to be further studied in AMR, together with metabolism.

**Figure 4.**
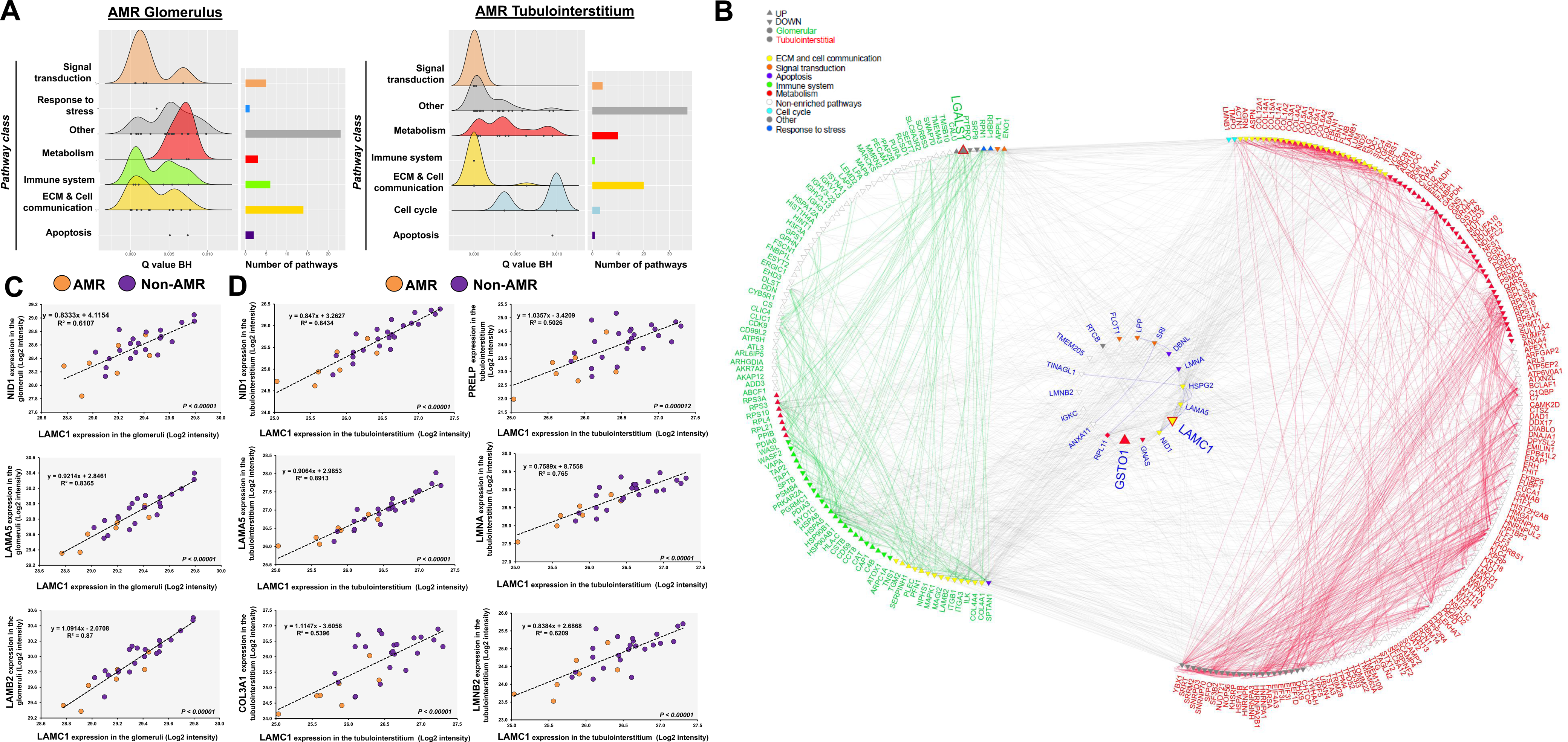
‘Extracellular Matrix’ represents the most enriched pathway category in both AMR compartments. For each compartment, the significantly enriched pathways from Pathdip were classified using a reduction algorithm. Pathway enrichment and reduction analysis revealed that the main category among the proteins significantly altered in the AMR vs. non-AMR glomeruli and tubulointerstitium was ‘ECM and cell communication’ (A). 300 proteins differentially expressed in one or both AMR compartments were analysed in the Integrated Integrations Database to obtain protein-protein interactions (PPIs), and only the ones annotated with ‘kidney’ were retained. The data was used to build a network in NAViGaTOR 3 (B). Each node represents a protein and, as shown in the legend, was colored according to one of the categories. Nodes belonging to non-significantly enriched pathways are shown in white. The orientation of the node vertex depicts the direction of change in AMR compared to non-AMR. Each edge represents a predicted or experimentally validated protein-protein interaction among proteins differentially expressed in the AMR vs. non-AMR glomeruli (green edges), tubulointerstitium (red edges), or both (blue edges) (B). Pearson correlation was calculated between the expression values of the 120 proteins differentially expressed in the AMR glomeruli (C), and between the expression values of the 180 proteins differentially expressed in the AMR tubulointerstitium (D). In both compartments, LAMC1 protein expression correlated directly, strongly, and significantly with the expression of other ECM proteins significantly downregulated in AMR.

Given the prominent role of LAMC1 in both AMR compartments, we verified the changes in LAMC1 expression, using immunohistochemistry. Consistent with the proteomics findings, numerically lower LAMC1 staining was detected in the AMR compared to non-AMR glomeruli and tubulointerstitium (**Fig.5A**). Since podocytes directly communicate with the glomerular BM, the impairment in podocyte-specific proteins identified in the AMR glomeruli may relate to the early ECM changes and glomerular BM-remodeling (**Fig.2**). Supporting our proteomics data, we observed reduced NPHS1 and PTPRO immunofluorescence, and a significant decrease in the merged signal of the two proteins, in AMR compared to non-AMR glomeruli (**Fig.5B**).

**Figure 5.**
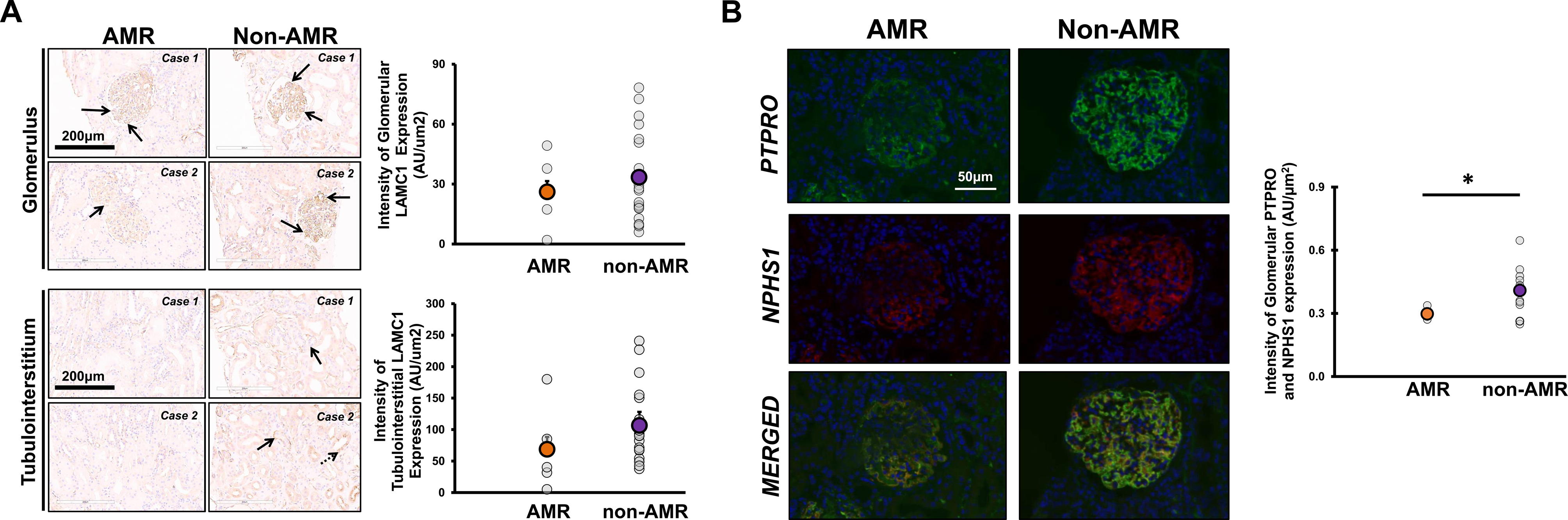
Orthogonal verification of representative basement membrane proteins. Decreased glomerular and tubulointerstitial protein levels of LAMC1 in AMR (n=6) vs. non-AMR (n=21) were verified by immunohistochemistry in new sections of formalin-fixed paraffin-embedded biopsies employed in the discovery study (A). Solid arrows indicate higher LAMC1 expression in the glomerular and tubular basement membranes of the non-AMR compared with AMR cases, whereas the doted arrow indicates strong cytoplasmic staining in non-AMR tubules. Magnification: 20x. Scale bar: 200µm. Significantly decreased glomerular protein expression of NPHS1 and PTPRO in AMR (n=3) vs. non-AMR (n=10) was demonstrated using immunofluorescence (B). Magnification: 40x. Scale bar: 50µm. *P<0.05 in AMR vs. non-AMR. AMR, antibody-mediated rejection; AU, arbitrary units; LAMC1, laminin subunit gamma-1; NPHS1, nephrin; PTPRO, receptor-type tyrosine-protein phosphatase.

### Galectin-1 and cathepsins are linked to ECM alterations in AMR

We investigated the potential causes of ECM-remodeling in each AMR compartment, considering two hypotheses: 1) Proteins decreased in AMR have decreased transcription; and 2) ECM-related proteins decreased in AMR are cleaved by proteases with increased expression/activity in AMR.

We took advantage of the biggest (to our knowledge) publicly-available transcriptomic dataset (GSE36059) comparing AMR biopsies to stable controls. We cross-referenced our proteins dysregulated in each AMR compartment with the 1,275 genes differentially expressed in AMR biopsies (Q<0.05 AMR vs. Stable control, **Table S8**). 16/20 proteins or genes significant in both datasets had a concordant direction of change, with the majority being significantly increased in AMR glomeruli (**Fig.6A**). HLA class I-mediated antigen presentation proteins (TAP1, TAP2) were among them, and also an important immunomodulatory protein, linked to ECM-remodeling, LGALS1^39, 40^ (**Fig.4B,6A**). Increased expression of several proteases (CTSS, CASP1, GZMA, and GZMB) was identified in GSE36059 (**Fig.6B**). Cathepsins and caspases can cleave ECM and the cytoskeleton and thus relate to our proteomics findings in AMR,^41–43^ whereas granzymes are mainly expressed by immune cells.^44, 45^

**Figure 6.**
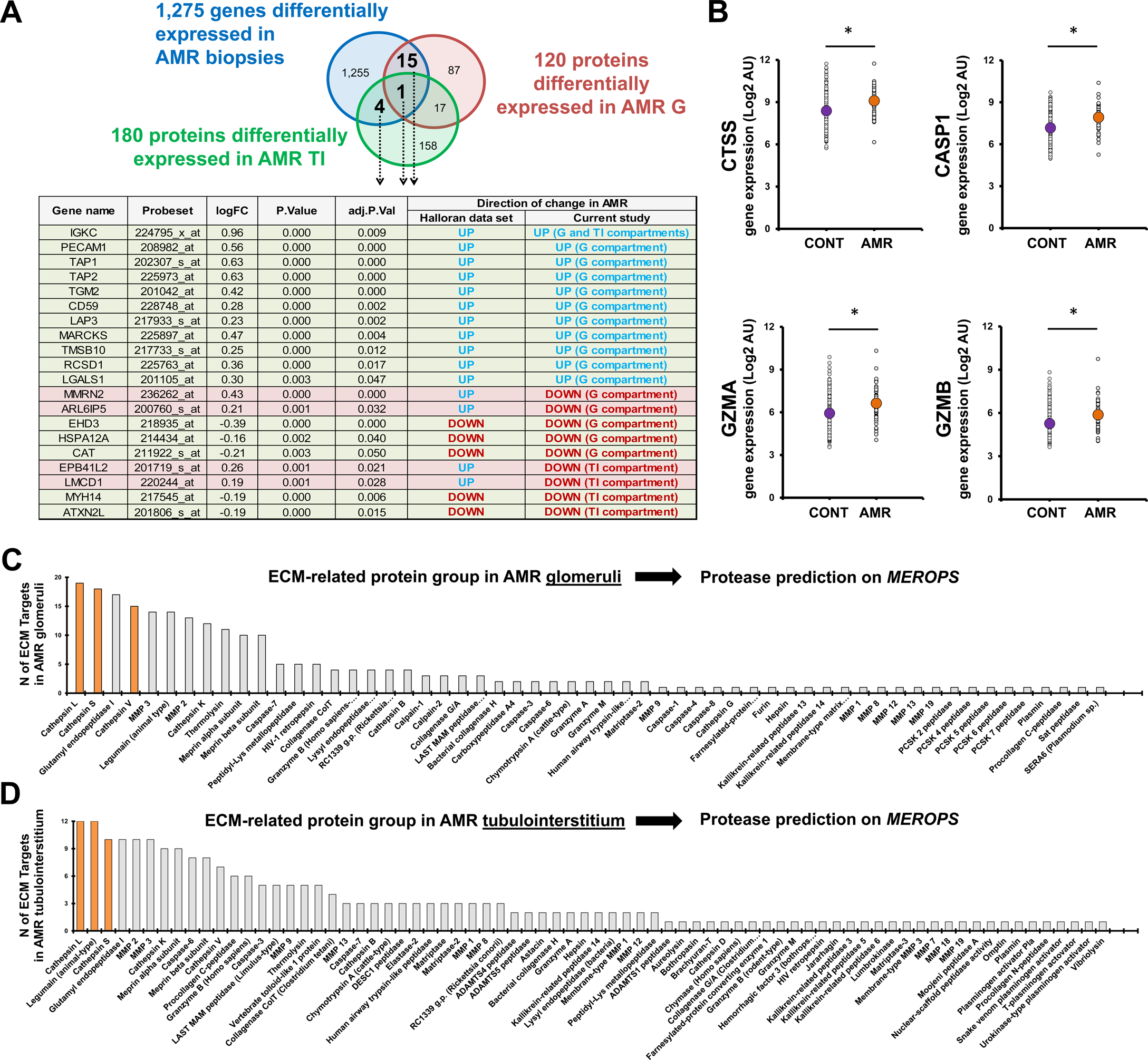
Comparison of our dataset to a relevant transcriptional dataset and protease cleavage prediction. Our findings at the proteome level were compared to the GSE36059 dataset. The Venn diagram in panel A represents the overlap between our three data sets of interest: 1) genes significantly differentially expressed in AMR kidney biopsies compared with control biopsies in GSE36059; 2) proteins significantly differentially expressed in the AMR glomeruli in our study; and 3) proteins significantly differentially expressed in the AMR tubulointerstitium in our study. As shown in the table, 20 proteins differentially expressed in AMR in our study were also dysregulated at the gene level in GSE36059. Sixteen of these 20 proteins changed in the same direction (green rows), and the remaining 4 changed in the opposite direction (red rows) between our and GSE36059 study. Proteases known to cleave ECM and cytoskeleton components, namely CTSS, CASP1, GZMA, and GZMB, were also significantly increased in GSE36059 (B). *Q<0.05 in AMR vs. control. ECM-related proteins dysregulated in each compartment in AMR were subjected to MEROPS protease prediction tool. The bar graphs represent the number of ECM-related proteins dysregulated (most of them downregulated) in the AMR glomeruli (C) or downregulated in the AMR tubulointerstitium (D) predicted to be cleaved by a specific protease. Top proteases predicted to cleave 10 or more ECM-related proteins are highlighted in orange. AMR, antibody-mediated rejection; G, glomeruli; TI, tubulointerstitium; CTSS, cathepsin S; CASP1, caspase-1; GZMA, granzyme A; GZMB, granzyme B; ECM, extracellular matrix.

We next used MEROPS^46^ to identify proteases predicted to cleave the ECM proteins in each AMR compartment. CTSL, CTSS, and CTSV were the top 3 proteases expressed in the kidney and predicted to cleave the highest number of ECM proteins dysregulated in the AMR glomeruli (**Fig.6C**). Together with legumain (LGMN), CTSL and CTSS were also predicted to cleave the highest number of ECM proteins decreased in the AMR tubulointerstitium (**Fig.6D**).

### Anti-HLA class-I antibodies increased galectin-1 in human microvascular glomerular endothelial cells

We studied the effects of anti-HLA class-I antibodies (α-HLA-I) and two prototypical cytokines associated with AMR, IFNɣ and TNFα, on our proteins and proteases of interest, in primary HGMECs. HGMECs exhibited the expected signaling, proliferative and inflammatory responses to α-HLA-I, IFNɣ and TNFα (**Fig.S4**). LGALS1 was predicted to interact with 14 other differentially expressed proteins in the AMR glomeruli, 7 of them in the ECM (**Fig.7A**). LGALS1 is at the nexus between metabolism, apoptosis, the immune system and ECM, and may represent a critical regulatory protein in AMR. Consistent with the proteomics and transcriptomics findings, α-HLA-I induced a significant increase in LGALS1 protein secretion and gene expression in HGMECs (**Fig.7B,C**). Similarly, both α-HLA-I and IFNγ significantly increased TAP1 gene expression (**Fig.7D**). TAP1 expression was increased in the AMR glomeruli (**Fig.2**) and in the transcriptomic dataset (GSE36059) (**Fig.6A**). In agreement with our protease prediction, stimulation of HMGECs with α-HLA-I but not isotype control, significantly increased CTSV gene expression, compared with vehicle-treated cells. IFNγ significantly enhanced CTSL, CTSS and CASP1 gene expression in HGMECs (**Fig.7E**). We assessed the expression of other proteases involved in ECM proteolysis, MMP2 and MMP9.^47, 48^ While α-HLA-I significantly upregulated MMP2 gene expression in HGMECs, IFNγ induced the opposite effect (**Fig.7E**).

**Figure 7.**
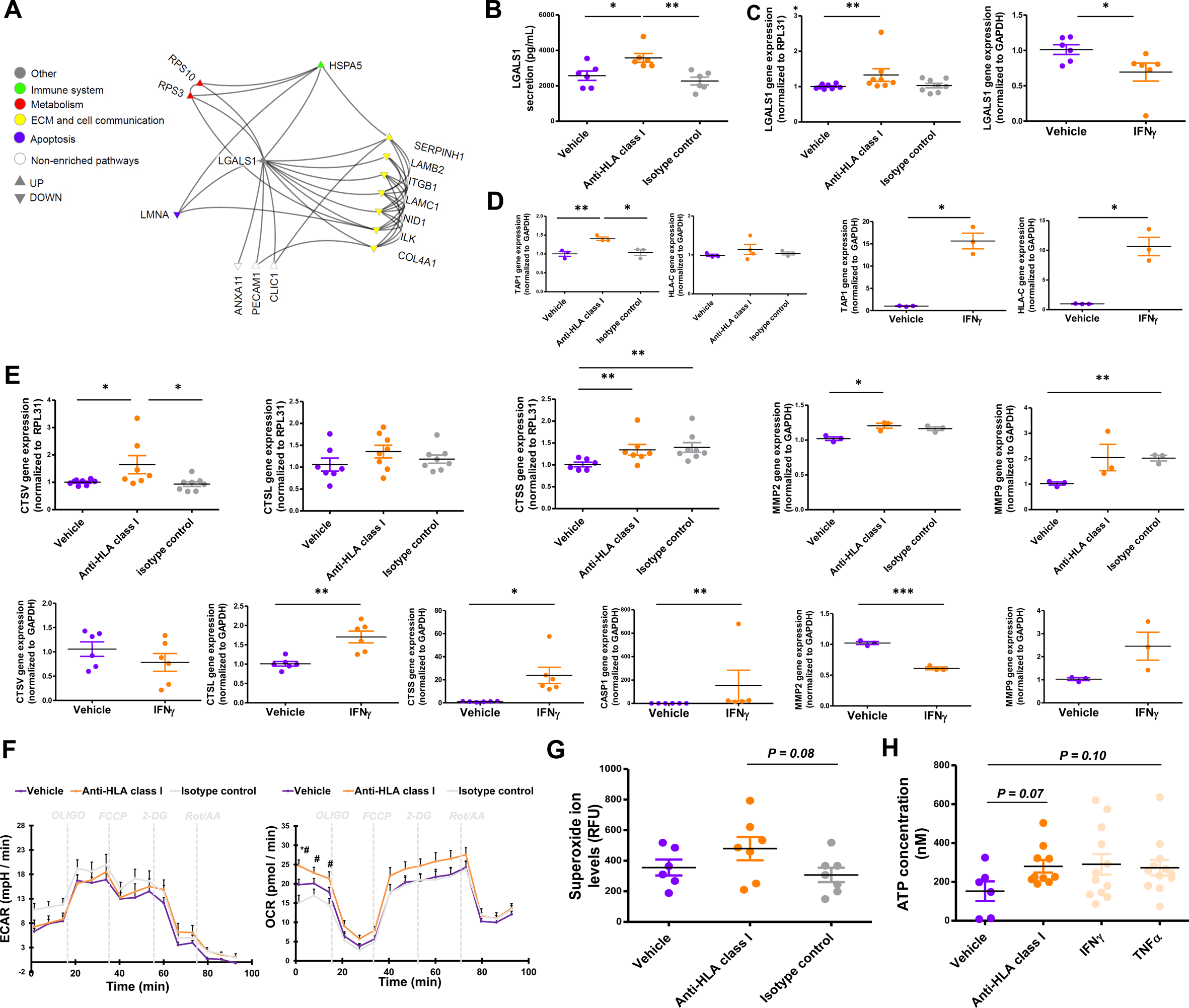
Validation of proteomic findings in the AMR glomeruli in primary human glomerular microvascular endothelial cells. Protein-protein interaction network of LGALS1 and its interactors among the 120 proteins differentially expressed in the AMR compared to non-AMR glomeruli (A). Only protein-protein interactions annotated in the kidney were retained, and the network was built in NAViGaTOR. Each node represents a protein, and each edge represents a predicted or experimentally validated protein-protein interaction. Each node was colored according to a specific pathway category. The orientation of the node vertex depicts the direction of change in AMR compared to non-AMR. Stimulation with 5µg/mL of α-HLA-I for 18h significantly upregulated LGALS1 protein secretion (B) and gene expression (C) in HGMECs. Gene expression of TAP1 and HLA-C in HGMECs was also upregulated by α-HLA-I and after stimulation with 1,000U/mL IFNγ for 24h (D). The effects of α-HLA-I and IFNγ on the gene expression of key proteases, namely CTSV, CTSL, CTSS, CASP1, MMP2, and MMP9, were also studied (E). Glycolysis (ECAR) and oxygen consumption rate (OCR) were monitored in a Seahorse XFe96 analyzer in HGMECs (F). The following concentrations were employed: oligomycin: 1µM; FCCP: 0.6µM, 2-DG: 100mM; Rot: 1mM; AA: 1mM. Oligomycin induces an increase in glycolysis by inhibiting ATP synthase. FCCP induces mitochondrial stress by uncoupling respiration from ATP synthesis. Rot/AA are electron transport chain inhibitors. *P<0.05 vs. Vehicle; #P<0.05 vs. isotype control-treated HGMECs. α-HLA-I stimulation in HGMECs increased levels of superoxide ion (G). Together with IFNγ and TNFα, α-HLA-I also induced an increase in the intracellular levels of ATP (H). Data are expressed as mean ± SEM. *P<0.05; **P<0.01; ***P<0.001. HGMECs, human glomerular microvascular endothelial cells; α-HLA-I, anti-HLA class I antibodies; IFNγ, interferon gamma; TNFα, tumor necrosis factor alpha; LGALS1, galectin-1; TAP1, antigen peptide transporter 1; HLA-C, HLA class I histocompatibility antigen, C alpha chain; CTSV, cathepsin-V; CTSL, cathepsin-L; CTSS, cathepsin-S; CASP1, caspase-1; MMP2, matrix metalloproteinase-2; MMP9, matrix metalloproteinase-9; GAPDH, glyceraldehyde-3-phosphate dehydrogenase; RPL31, 60S ribosomal protein L31; ECAR, extracellular acidification rate; OCR, oxygen consumption rate; FCCP, p-trifluoromethoxy carbonyl cyanide phenyl hydrazone; 2-DG, 2-deoxyglucose; Rot, rotenone; AA: antimycin A.

Since metabolism emerged as an important process enriched in the AMR glomeruli, we examined the effects of α-HLA-I on the HGMEC metabolic function. Without affecting glycolysis, α-HLA-I antibodies induced a significant increase in the oxygen consumption rate (**Fig.7F**) and numerically increased cellular levels of superoxide ion (**Fig. 7G**) and ATP (**Fig. 7H**) in HGMECs. ATP was also numerically increased by IFNγ and TNFα.

### TNFα increased GSTO1 expression in human proximal tubular epithelial cells

TNFα and IFNɣ-mediated pathways were enriched in the AMR tubulointerstitium (**Table S5**). We thus stimulated PTECs with IFNɣ and TNFα, and demonstrated the expected proinflammatory response to these cytokines (**Fig.S5**). We next focused on GSTO1, a metabolic protein increased in the AMR tubulointerstitium (**Fig.4B**) that modifies ECM proteins and increases their susceptibility to proteolytic cleavage^49^, representing a potential link between metabolism and ECM-related alterations. TNFα significantly increased GSTO1 protein expression in PTECs (**Fig.8A**). In line with our protease prediction, TNFα significantly increased CTSS expression, compared to vehicle, in PTECs. This increase was more pronounced upon stimulation with IFNɣ, which also dramatically increased LGMN (**Fig.8B**).

**Figure 8.**
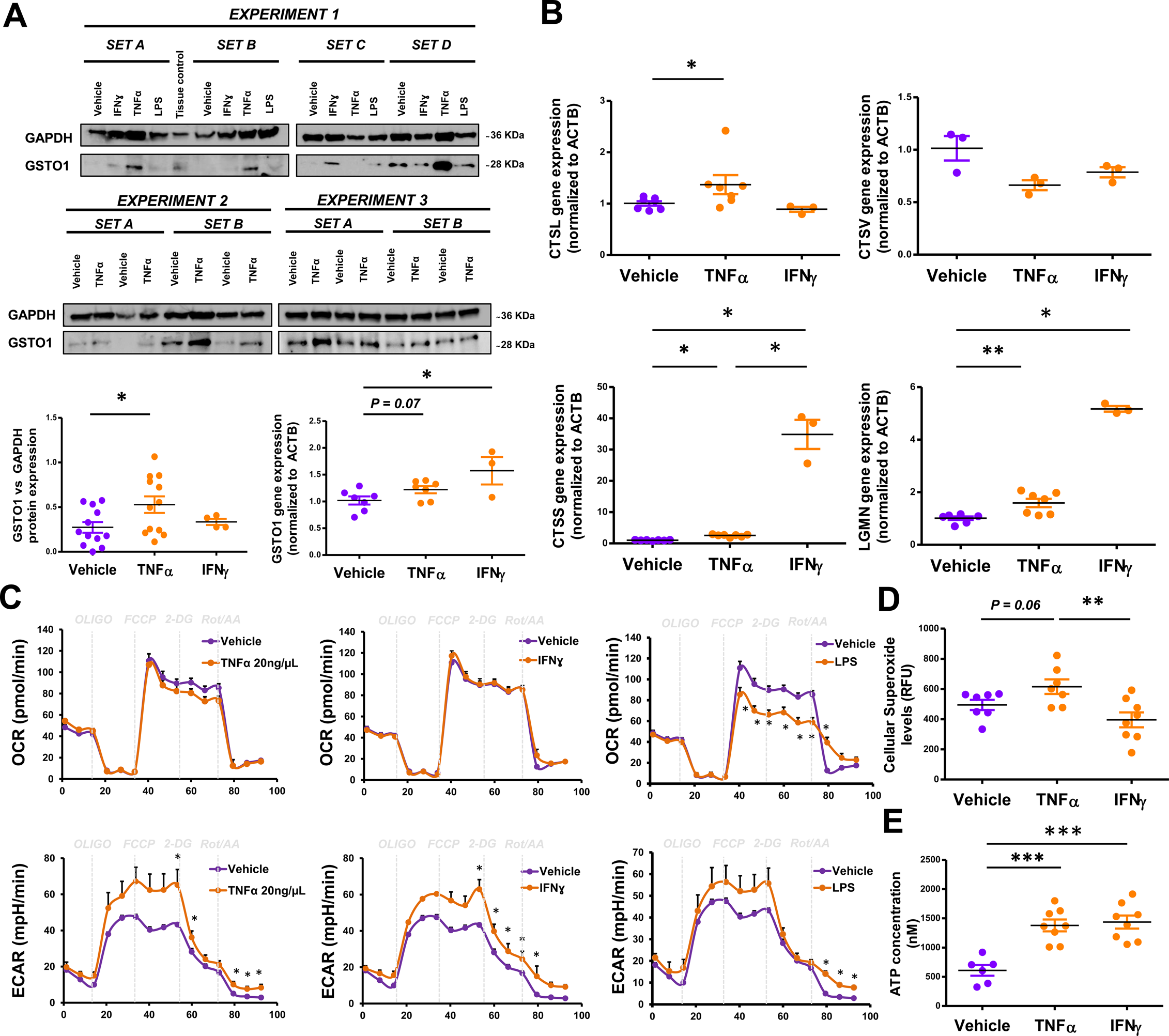
Validation of proteomic findings in the AMR tubulointerstitium in primary human proximal tubular epithelial cells Protein expression of GSTO1 was analyzed in PTECs after treatment with vehicle, 20ng/µL of TNFα, 1000U/mL of IFNγ, or 1ng/µL of LPS for 24h, and normalized to GAPDH (A). The immunoblots and corresponding densitometry values are shown. GSTO1 expression was also measured at the gene level after treatment with vehicle, 20ng/µL of TNFα, or 1000U/mL of IFNγ for 24h (A). The effects of TNFα and IFNγ on the gene expression of CTSL, CTSV, CTSS, and LGMN proteases, were studied (B). OCR and ECAR were monitored in a Seahorse XFe96 analyzer in order to study the effects of TNFα, IFNγ, or LPS on PTECs. As expected, LPS-treated PTECs (positive control) experienced a metabolic switch towards a more glycolytic phenotype, compared to vehicle-treated cells (C). The following concentrations were employed: oligomycin: 1µM; FCCP: 0.3µM, 2-DG: 100mM; Rot: 1mM; AA: 1mM. Oligomycin induces an increase in glycolysis by inhibiting ATP synthase. FCCP induces mitochondrial stress by uncoupling respiration from ATP synthesis. Rot/AA are electron transport chain inhibitors. *P<0.05 vs. Vehicle-treated PTECs. Intracellular levels of superoxide ion (D) and ATP (E) were also measured in vehicle-, TNFα-, and IFNγ-treated PTECs. Data are expressed as mean ± SEM. *P<0.05; **P<0.01; ***P<0.001. PTECs, human proximal tubular epithelial cells; TNFα, tumor necrosis factor alpha; IFNγ, interferon gamma; LPS, lipopolysaccharide; GSTO1, glutathione S-transferase omega-1; GAPDH, glyceraldehyde-3-phosphate dehydrogenase; CTSL, cathepsin-L; CTSV, cathepsin-V; CTSS, cathepsin-S; LGMN, legumain; ACTB, beta-actin; ECAR, extracellular acidification rate; OCR, oxygen consumption rate; FCCP, p-trifluoromethoxy carbonyl cyanide phenyl hydrazone; 2-DG, 2-deoxyglucose; Rot, rotenone; AA: antimycin A.

Since GSTO1 is directly involved in metabolism and the redox balance of the cell, we studied the metabolic function of cytokine-treated PTECs. Both TNFα and IFNɣ increased glycolysis in PTECs, especially after metabolic stress (**Fig.8C**). Although TNFα and IFNɣ did not significantly alter oxygen consumption or superoxide ion levels (**Fig.8C,D**), both cytokines induced a significant increase in intracellular ATP (**Fig.8E**). PTECs thus display a higher dependence on glycolysis for energy production, upon TNFα and IFNɣ stimulation.

## DISCUSSION

This is the first proteomics study of laser-captured/microdissected glomeruli and tubulointerstitium in kidney AMR biopsies compared to other forms of graft injury. Our main observation is that AMR is associated with a compartment-specific decrease in ECM-protein expression, before histological signs of BM injury or fibrosis. This ECM-remodeling was not observed in transcriptional studies;^27, 50–53^ instead, our work suggests that it is associated with dysregulated protease expression. LGALS1 and GSTO1 were upregulated in the glomerular and tubulointerstitial AMR proteomes respectively, and validated in models of anti-HLA and TNFα-mediated injury. These ECM-modifying proteins may represent novel targets to ameliorate ECM-remodeling associated with AMR.

Importantly, ECM proteins downregulated in the AMR glomeruli (nidogen, collagen-IV chains, and laminin subunits) belong to the glomerular BM,^54^ which is critical to the integrity of the filtration barrier.^55^ Double-contouring of BMs is a hallmark of TG.^56, 57^ Our AMR cases did not show such injury histologically. Protein changes in AMR glomeruli could thus reflect an ECM-remodeling starting early in AMR. Glomerular BM is secreted by endothelial cells and podocytes, representing an interface between them.^58^ Glomerular BM proteins anchor podocytes via adhesion receptors connected to the cytoskeleton.^54^ Several adhesion proteins were downregulated in AMR. Thus, the interactions between endothelium, BM and podocytes may be compromised in early AMR. We observed a significant downregulation of slit diaphragm proteins, NPHS1 and PTPRO, in AMR.^59, 60^ PTPRO-deficient mice developed abnormal podocytes, but also remodeled glomerular BM, suggesting that PTPRO is important for both podocyte and glomerular BM integrity.^61, 62^ Reduced PTPRO expression correlated with podocyte loss in TG.^63^ Although we did not detect podocyte effacement, decreased BMs proteins occurred in association with altered integrity of podocyte proteins in AMR.

Graft endothelium is central to antibody-mediated injury, and we next focused on linking endothelial cell injury to ECM-remodeling. LGALS1 was particularly interesting, because it was increased in our glomerular AMR proteome and the AMR transcriptome,^27^ and it is secreted by endothelial cells,^40^ and linked to ECM-remodeling.^39^ LGALS1 recognizes β-galactose moieties of ECM proteins,^39^ and is involved in immunomodulation, angiogenesis, survival and proliferation.^64, 65^ We demonstrated that α-HLA-I increased LGALS1 in HGMECs. Similar to previous studies, this was related to increased cell proliferation, signaling and inflammation.^14, 15, 66^ A protective role for LGALS1 was previously proposed. LGALS1^-/-^ animals displayed enhanced inflammation and oxidative stress,^67^ and transfer of B cells from LGALS1^-/-^ mice failed to prolong skin allograft survival in mice.^67^ Conversely, recombinant LGALS1 decreased inflammation in renal ischemia-reperfusion injury model,^68^ and prolonged graft survival in a model of MHC-mismatched kidney transplantation.^69^ LGALS1 may thus participate in the graft endothelial response to DSA.

This is the first study that investigates the tubulointerstitial proteome in AMR. We observed significantly decreased collagens and BM proteoglycans in the AMR tubulointerstitium. Tubular BM disruption may cause abnormal tubular cell function and tissue destruction.^70^ We thus studied ECM-modifying enzymes in PTECs exposed to TNFα and IFNɣ. GSTO1 was upregulated in the AMR tubulointerstitium and in TNFα-treated PTECs. GSTO1 is an enzyme that mediates S-deglutathionylation of ECM and cytoskeletal proteins, increasing the pool of cytosolic glutathione capable of neutralizing reactive oxygen species.^71^ However, S-deglutathionylation also affects the ECM-cytoskeleton network, altering protein susceptibility to proteolytic cleavage.^49^ TNFα-induced GSTO1 upregulation in PTECs was linked to augmented glycolysis, ATP levels, oxidative stress, and inflammation. Cytokine-activated immune cells rely on glycolysis for ATP generation, to support their pro-inflammatory phenotype.^72^ In turn, TNFα impairs mitochondrial function in renal tubular and other cells, probably forcing them to use glycolysis for ATP production.^73, 74^ Interestingly, an inflammatory role of GSTO1 has been described in macrophages.^75^ Thus, increased GSTO1 in AMR may reflect a maladaptive response to TNFα-induced metabolic stress.

Altered ECM protein expression in AMR could be attributed to increased proteolytic cleavage (although intracellular degradation was not explored). Cathepsins degrade ECM proteins, including collagens, laminins, and proteoglycans.^76–79^ Additionally, NID1 binding to the BM is impaired after cleavage by CTSS,^80^ which was transcriptionally enhanced in kidney AMR^27^ and in response to IFNɣ^81–83^. In PTECs, CTSS and LGMN were upregulated by TNFα and IFNɣ. LGMN inhibition reduced IFNγ production, CTSL activation and C3a generation in activated human CD4^+^ T-cells; moreover, CD4+ T-cells from *Lgmn*-deficient mice showed defective IFNγ production and Th1 induction.^84^ CTSS and LGMN also mediate HLA class-II antigen presentation, promoting the generation of competent HLA class-II molecules for peptide binding,^85, 86^ and processing of exogenous peptides for HLA class-II-presentation,^87–89^ respectively. CTSS inhibition prevented HLA class-II-mediated antigen presentation in autoimmunity,^90^ and reduced the generation of competent HLA class-II and the infiltration of IFNɣ-expressing CD4+ T-cells in a neuropathy model.^91^ Cathepsins also interact with LGALS1.

Accordingly, increased IL6, IL8, and LGALS1 in α-HLA-I-treated HGMECs coincided with increased CTSV expression. CTSV mediates inflammation in HUVECs by upregulating these cytokines.^92^ LGALS1 binds to membrane glycoproteins, inhibiting their endocytosis and cathepsin-mediated cleavage,^93^ while cathepsins induce LGALS1 in endothelial cells during angiogenesis.^94^ Whether decreased expression of cathepsin targets in AMR is due to higher cleavage and higher HLA-presentation requires further investigation.

This work has limitations. The number of cases was small, as we focused on extreme phenotypes, and were constrained by cases with predominantly mixed lesions or limited available tissue. We focused on injury mechanisms in kidney parenchymal cells, but acknowledge that immune cells also produce ECM-modulating proteins. Cell models cannot recapitulate the complexity of *in vivo* systems. Future studies will determine the relationship between DSA and BM-remodeling *ex vivo* and *in vivo*.

We have identified compartment-specific ECM-remodeling in the absence of BM lesions, in early AMR. We propose a model of antibody- and cytokine-mediated injury in the AMR glomeruli and tubulointerstitium (**Fig.9**). Decreased BM and ECM proteins in the AMR glomeruli were associated with α-HLA-I-induced upregulation of antigen-presenting (TAP1) and ECM-modifying proteins (LGALS1 and CTSV), and with IFNɣ-induced upregulation of key proteases (CTSL, CTSS). In the AMR tubulointerstitium, decreased ECM-related proteins were associated with TNFα- and IFNɣ-induced upregulation of ECM-modifying enzymes (GSTO1, CTSS, CTSL and LGMN). Our findings thus point to early, potentially targetable alterations in AMR.

**Figure 9.**
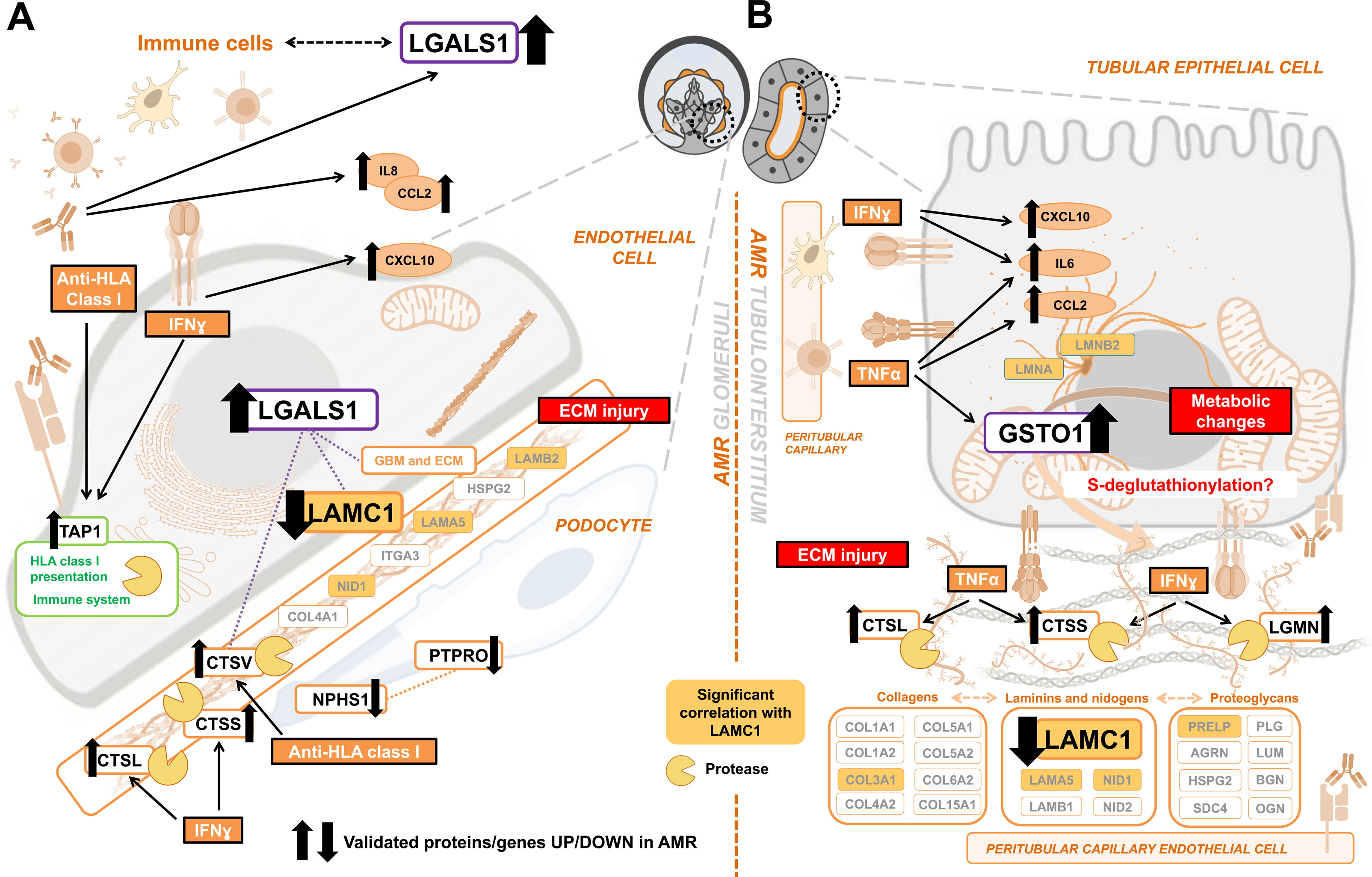
Proposed model of cell-specific extracellular matrix injury in AMR. Simplified scheme depicting the key proteins and processes differentially expressed in the AMR vs. non-AMR glomeruli (A) and tubulointerstitium (B). Protein name boxes are colored according to the category they belong to. The direction of change is indicated for the proteins verified by immunohistochemistry or validated *in vitro.* Relevant protein-protein interactions between LGALS1, LAMC1, cathepsins, and AMR-enriched processes are represented by dotted lines. Proposed biological processes and molecular alterations associated to AMR are summarized in red boxes. AMR, antibody-mediated rejection; ECM, extracellular matrix; GBM, glomerular basement membrane; HLA, human leukocyte antigen; ER, endoplasmic reticulum; TNFα, tumor necrosis factor alpha; IFNγ, interferon gamma; LAMC1, laminin subunit gamma-1; LGALS1, galectin-1; NPHS1, nephrin; PTPRO, receptor-type tyrosine-protein phosphatase O; TAP1, antigen peptide transporter 1; GSTO1, glutathione S-transferase omega-1; CTSL, cathepsin-L; CTSV, cathepsin-V; CTSS, cathepsin-S; LGMN, legumain; CCL2, C-C motif chemokine 2; CXCL10, C-X-C motif chemokine 10; IL6, interleukin-6; IL8, interleukin-8.

## AUTHOR CONTRIBUTIONS

AK conceived the study. AK and RJ participated in study design; SC-F, CMcE, IB, SF, JV, SM, MA, Alex B, AC, Andrea B, VK, RJ, and AK carried out experiments; SC-F, CMcE, IB, MK, CP, YN, IJ, SM, SJ, MA, TM, RJ, and AK analyzed the data; SC-F, CMcE, CP, and MK made the figures; CMcE, YL, PC, EC, retrieved and curated clinical data; YL, OF and SJK selected the cases from the CoReTRIS registry and performed case-control matching; SC-F, CMcE, and AK drafted and revised the paper; all authors approved the final version of the manuscript.

## ACKNOWLEDGEMENTS

Special thanks to Marc Angeli, Sharon Selvanayagam, Terrance Ku, Victor Ferreira, Bedra Sharif, Matthew Ierullo, Beata Majchrzak-Kita and Elisa Pasini.

## Funding

AK is supported by a Kidney Foundation of Canada operating grant, the Kidney Research Scientist Core Education and National Training (KRESCENT) program, Kidney Foundation of Canada Predictive Biomarker Grant, Canadian Institute for Health Research (CIHR), and Canada Foundation for Innovation (CFI). She has also received funding from the Toronto General Hospital Research Foundation and the Multi-Organ Transplant program. SC-F is supported by the KRESCENT program and CIHR Catalyst Grant. CMcE is supported by the Menkes fellowship, and a University Health Network Multi-Organ Transplant fellowship. IJ, CP & MK were supported in part by Ontario Research Fund (#34876), Natural Sciences Research Council (NSERC #203475), CFI (#29272, #225404, #30865), Krembil Foundation and IBM.

## DISCLOSURES

Nothing to disclose.

## SUPPLEMENTAL MATERIAL - TABLE OF CONTENTS

## FIGURE LEGENDS – SUPPLEMENTAL

## Abbreviations

ACR: Acute cellular rejection
AMR: Antibody-mediated rejection
ATN: Acute tubular necrosis
BM: Basement membrane
CTSL: Cathepsin-L
CTSS: Cathepsin-S
CTSV: Cathepsin-V
DSA: Donor-specific antibodies
ECM: Extracellular matrix
ELISA: Enzyme-linked immunosorbent assay
FDR: False discovery rate
GSTO1: Glutathione S-transferase omega-1
HGMEC: Human glomerular microvascular endothelial cells
HLA: Human leucocyte antigen
IFNɣ: Interferon gamma
LAMC1: Laminin subunit gamma-1
LFQ: Label-free quantification
LGALS1: Galectin-1
LGMN: Legumain
MS/MS: Tandem mass spectrometry
NPHS1: Nephrin
PTEC: Proximal tubular epithelial cells
PTPRO: Receptor-type tyrosine-protein phosphatase O
TG: Transplant glomerulopathy
TNFα: Tumor necrosis factor-alpha

---

**Figure S1.** Histograms depicting the distribution of the original and imputed protein intensity values in our study samples. Each histogram represents the distribution of the log2 transformed intensity values among the proteins quantified in each of the biopsy samples in the glomeruli (A, n=28) or tubulointerstitium (B, n=30). Blue bars represent the count of intensity values determined by mass spectrometry, whereas red bars represent the distribution of the imputed intensity values. Two of the glomerular fractions were excluded from further analyses due to poor protein recovery, which was reflected in a flat, non-normal distribution.

**Figure S2.** Representative light and electron microscopic images and injury scores of ultrastructural alterations in AMR and non-AMR cases. Representative light microscopy images of AMR, ACR and ATN cases (A, 20X). Representative electron microscopy images of a glomerular capillary loop in each study group (B, 8000X). Bar graphs showing glomerular basement membrane thickness (C), proportion of loops with evidence of subendothelial new basement membrane formation (relative to the total number of loops studied in two glomeruli per biopsy (D)), and scores (0-3) of glomerular (E) and peritubular (F) endothelial cell swelling (4 to 7 cases/group were studied). Semiquantitative score legend: 0 = none; 1 = mild (<25%); 2 = moderate (25-50%); 3 = severe (>50%). AMR, antibody-mediated rejection; ACR, acute cellular rejection; ATN, acute tubular necrosis; GBM, glomerular basement membrane.

**Figure S3.** Principal component analysis of the differentially expressed proteins in the AMR glomeruli and tubulointerstitium. Principal component analysis (PCA) of the proteins differentially expressed in AMR in each compartment (120 in the glomeruli (A) and 180 in the tubulointerstitium (B)) was performed and plotted using ‘ggbiplot’ (Version 0.55) and RStudio (Version 1.1.463) in R (Version 3.6.1)^31^. Twenty-eight samples were analyzed in the glomerular compartment (7 AMR, 10 ACR, and 11 ATN cases) and 30 in the tubulointerstitial compartment (7 AMR, 11 ACR, and 12 ATN cases). For each sample, the log2-transformed LFQ intensity value of each protein was used for the PCA. AMR, antibody-mediated rejection; ACR, acute cellular rejection; ATN, acute tubular necrosis.

**Figure S4.** Characterization of human glomerular microvascular endothelial cells. To confirm that HGMECs exhibited the expected endothelial cell phenotype at passage 5, the expression of endothelial cell-specific markers PECAM1, CHD5, and VWF in HGMECs, but not in other cell lines (namely human lung fibroblasts and renal proximal tubular epithelial cells (PTECs) was corroborated by real-time quantitative PCR and normalized to GAPDH. Lower expression of ACTA2 in HGMECs as compared to fibroblasts was also confirmed (A). HGMECs displayed the expected change in cell morphology towards a more oval, spindle-shaped phenotype in response to treatment with 1000U/mL IFNɣ or 10ng/mL TNFα for 24h (B). Magnification: 20x. Scale bar: 20µm. The phenotype changes after cytokine treatment were accompanied by a significant increase in CXCL10 and ACTA2 gene expression (C). Western blot and subsequent densitometry analysis showed that HGMECs expressed HLA class I but not HLA class II protein at baseline. Moreover, treatment with 500U/mL or 1000U/mL IFNɣ for 24h induced HLA II protein expression, and 1000U/mL IFNɣ increased HLA I protein expression, which was most accentuated 48 hrs after treatment (D). Both vehicle- and IFNɣ-pretreated HGMECs displayed the expected rapid increase in Erk phosphorylation upon stimulation with 1µg/mL α-HLA-I (E). Stimulation of HGMECs with 1µg/mL α-HLA-I, 1000U/mL IFNɣ or 10ng/mL TNFα for 24h elicited a proliferative response, as evidenced by increased intracellular RNA and DNA levels (F). To characterize the inflammatory response of HGMECs to α-HLA-I stimulation, cells were exposed to vehicle or 1µg/mL α-HLA-I for 24h, and secreted levels of IL-8, CCL2, and IL-4 were assessed in the cell supernatant by Multiplex ELISA (G). Gene expression levels of IL8, IL6, and VCAM1 (G) were also measured in HGMECs after exposure to vehicle, 5µg/mL of α-HLA-I, or 5µg/mL of isotype control for 12h, and normalized to VCL (H). Data are represented as mean ± SEM. *P<0.05; **P<0.01; ***P<0.001. HGMECs, human glomerular microvascular endothelial cells; PTECs, proximal tubular epithelial cells; ACTA2, aortic smooth muscle actin; PECAM1, platelet endothelial cell adhesion molecule; CHD5, cadherin-5; vWF, von Willebrand factor; GAPDH, glyceraldehyde-3-phosphate dehydrogenase; CXCL10, C-X-C motif chemokine 10; MHC, major histocompatibility complex; IFNγ, interferon gamma; TNFα, tumor necrosis factor alpha; α-HLA-I, anti-HLA class I antibodies; ERK, mitogen-activated protein kinase; IL-4, interleukin-4; IL6, interleukin-6; CXCL8/IL-8, interleukin-8; CCL2, C-C motif chemokine 2; VCAM1, vascular cell adhesion protein 1; VCL, vinculin.

**Figure S5.** Characterization of the inflammatory response of PTECs upon exposure to key cytokines in the AMR tubulointerstitium PTECs displayed a modest change in cell morphology towards a more oval, spindle-shaped phenotype in response to treatment with 20ng/mL TNFα or 1ng/mL LPS for 24h. This change was more evident upon treatment with 1000U/mL IFNɣ for 24h (A). Magnification: 20x. Scale bar: 50µm. To relate morphological changes to the expected dysregulation of proinflammatory genes after cytokine treatment, the gene expression of IL6 (B), CCL2 (C), and CXCL10 (D) was determined and normalized to ACTB. Data are represented as mean ± SEM. *P<0.05; **P<0.01; ***P<0.001. PTECs, proximal tubular epithelial cells; IL6, interleukin-6; CCL2, C-C motif chemokine 2; CXCL10, C-X-C motif chemokine 10; ACTB, beta-actin; IFNγ, interferon gamma; TNFα, tumor necrosis factor alpha; LPS, lipopolysaccharide.

**Table S1.** Primer sequences used for real-time quantitative PCR in our gene expression studies Table S2. Quantified and differentially expressed proteins in the AMR glomeruli and tubulointerstitium, compared to non-AMR

**Table S3.** Most representative gene ontology terms and pathways significantly enriched among the proteins differentially expressed in the AMR glomeruli, compared to non-AMR

**Table S4.** Most representative gene ontology terms and pathways significantly enriched among the proteins differentially expressed in the AMR tubulointerstitium, compared to non-AMR

**Table S5.** Protein interactors, and pathways significantly enriched among them, of the proteins upregulated in the AMR glomeruli and tubulointerstitium, compared to non-AMR

**Table S6.** Mapping between pathways significantly enriched in AMR compared to non-AMR and their ancestors

**Table S7.** Heatmap depicting the Pearson correlation coefficients between the log2 transformed LFQ intensity values of all the proteins significantly regulated in the AMR glomeruli and tubulointerstitium, compared to non-AMR

**Table S8.** Gene expression changes between AMR (n=65) and control (n=281) kidney biopsies (publicly available data set with identifier GSE36059)

