## Supplementary figures and images for "Proteomics Reveals Extracellular Matrix Injury in the Glomeruli and Tubulointerstitium of Kidney Allografts with Early Antibody-Mediated Rejection"

### Supplemental Figures

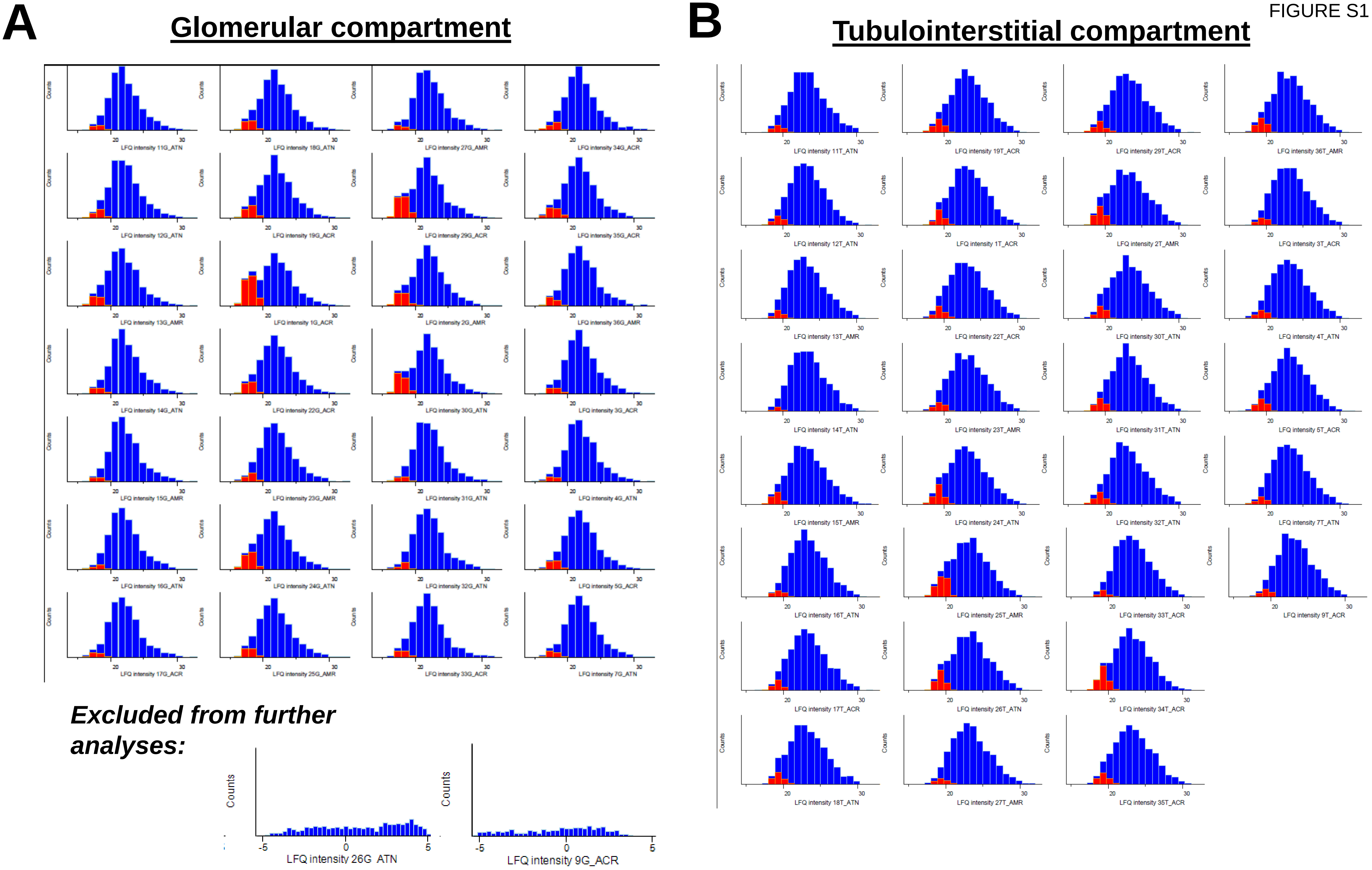

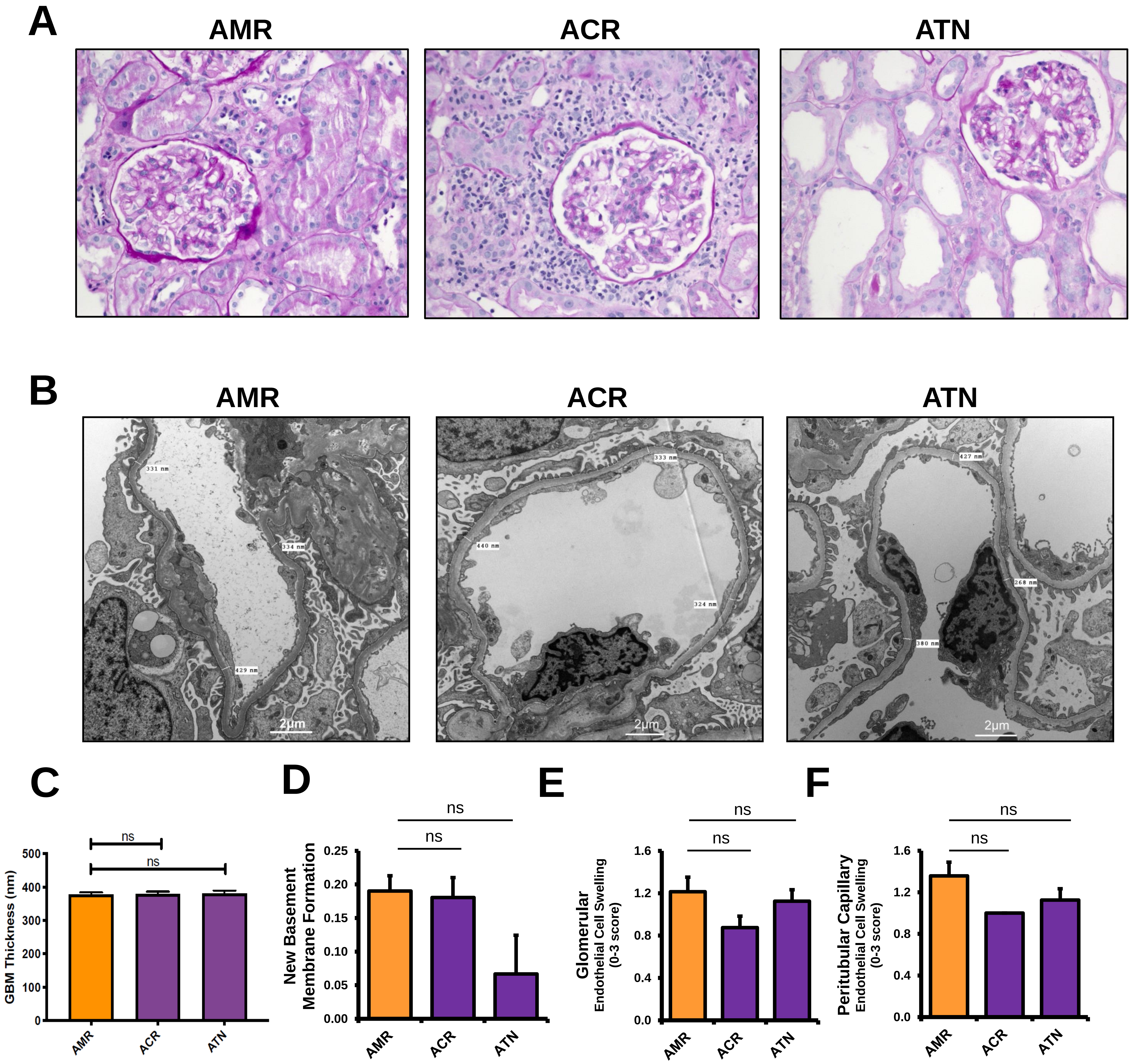

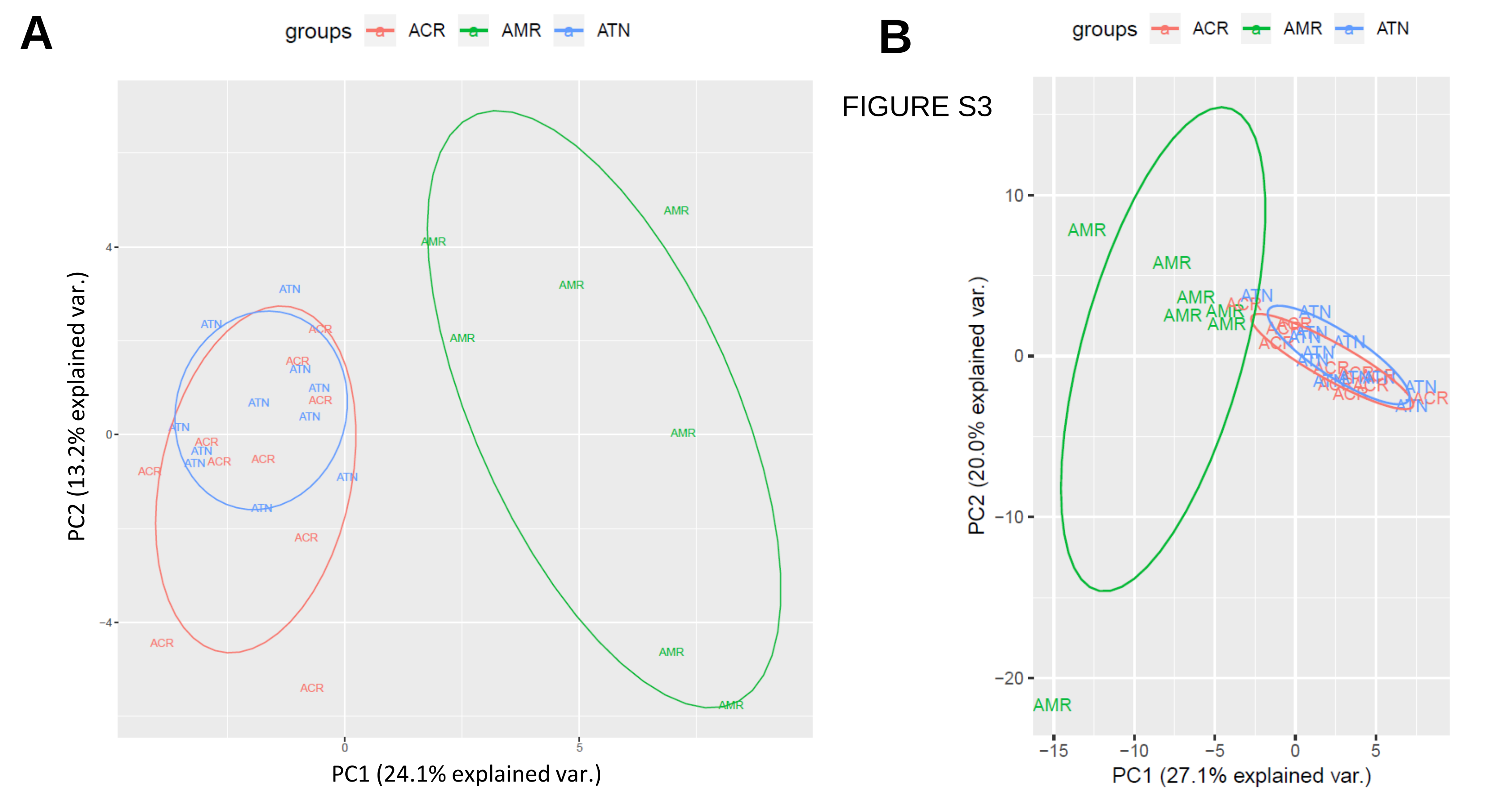

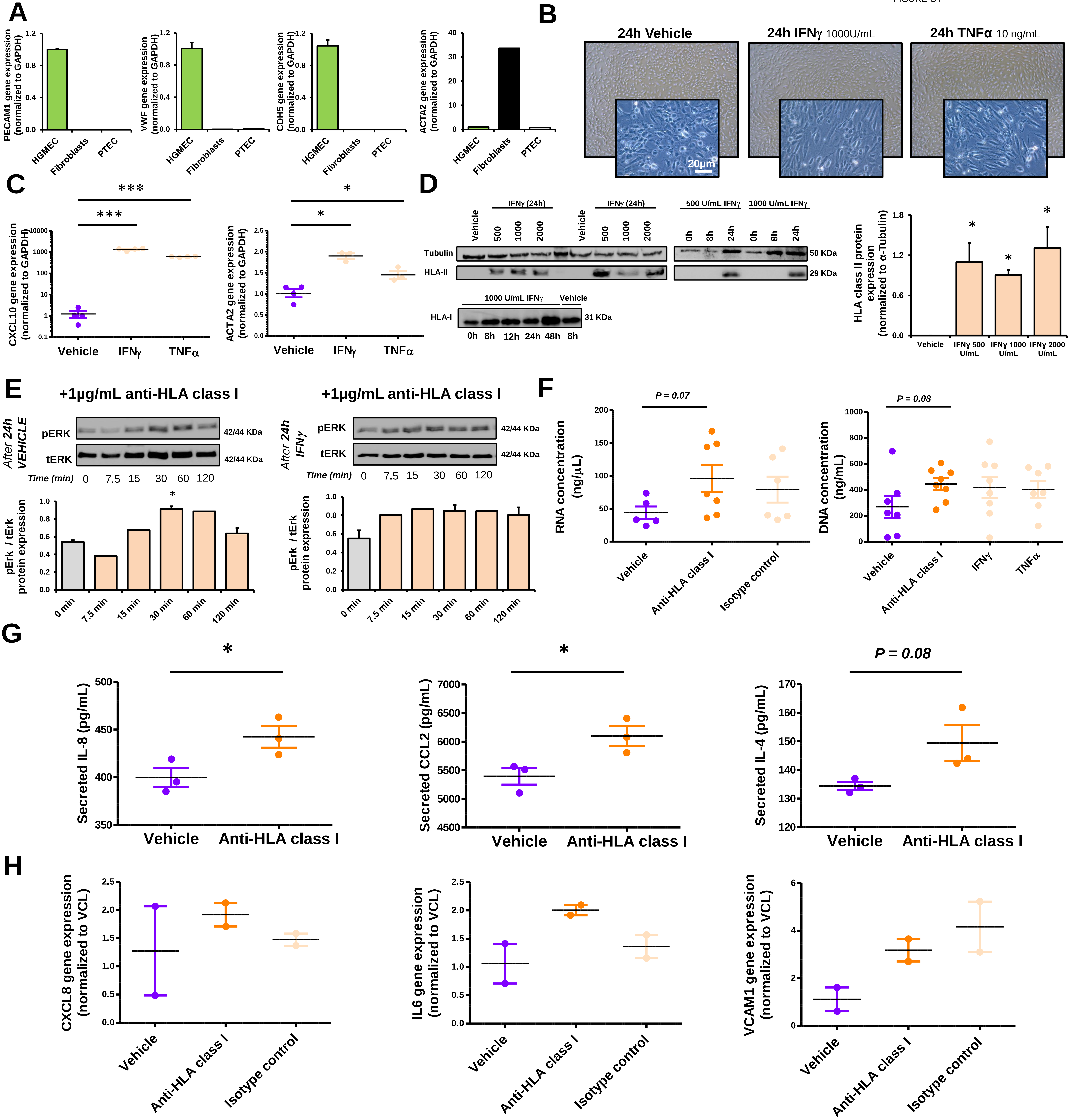

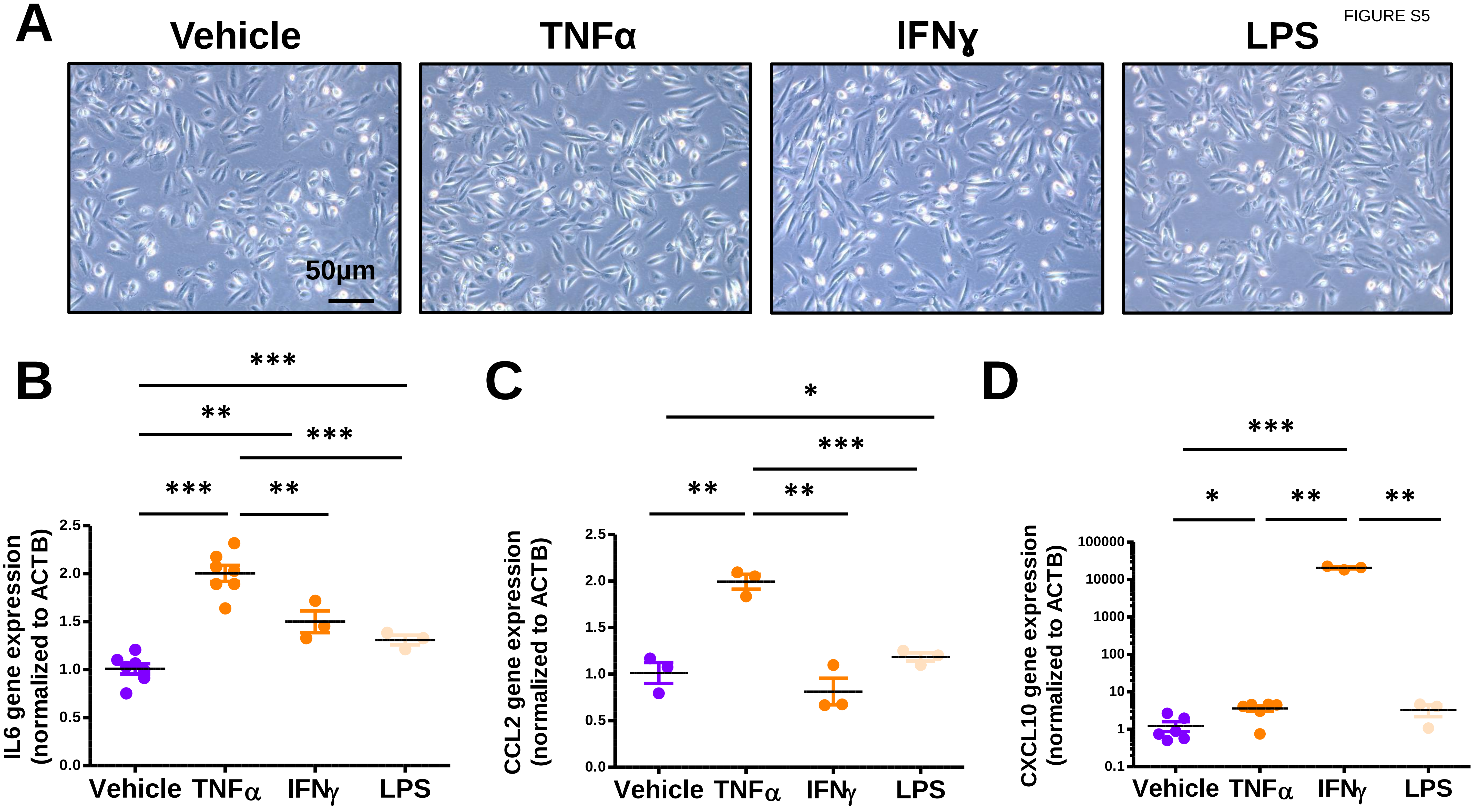
